# Education and income show heterogeneous relationships to lifespan brain and cognitive differences across European and US cohorts

**DOI:** 10.1101/2020.10.12.335687

**Authors:** Kristine B. Walhovd, Anders M. Fjell, Yunpeng Wang, Inge K. Amlien, Athanasia M. Mowinckel, Ulman Lindenberger, Sandra Düzel, David Bartrés-Faz, Klaus P. Ebmeier, Christian A Drevon, William Baaré, Paolo Ghisletta, Louise Baruël Johansen, Rogier A. Kievit, Richard N. Henson, Kathrine Skak Madsen, Lars Nyberg, Jennifer Harris, Cristina Solé-Padullés, Sara Pudas, Øystein Sørensen, René Westerhausen, Enikő Zsoldos, Laura Nawijn, Torkild Hovde Lyngstad, Sana Suri, Brenda Penninx, Ole J. Røgeberg, Andreas M. Brandmaier

## Abstract

Socio-economic status (SES) has been proposed to have facilitating and protective effects on brain and cognition. Here we show that relationships between SES, brain volumes and general cognitive ability differ significantly across European and US cohorts (4-97 years, N ≈ 500,000; 54,000 with brain imaging). Education was positively related to intracranial (ICV) and total brain gray matter (GM) volume. Income was related to ICV, but not GM. Relationships varied significantly across samples, and SES was more strongly related to brain and cognition in US than European cohorts. Differences in neuroanatomical volumes explained part of the SES-cognition relationships. SES was more strongly related to ICV than to GM, implying that SES-cognition relations in adulthood are less likely grounded in neuroprotective effects on GM volume in aging. Rather, a relationship may be established early in life. The findings underscore that SES has no uniform association with, or impact on, brain and cognition.

## Main

Socio-economic status (SES) has been proposed to have facilitating and protective effects on brain and cognition ^1–3^, and has been used as a proxy for cognitive reserve over the lifespan ^4^. Positive relationships between education, income, general cognitive ability (GCA) and brain volumes have been reported in development, adulthood and aging ^1,2,5–10^. SES variables are also frequently used as covariates of no interest. However, SES variables may not have a unified meaning or relation to brain and cognition across cohorts of varying ages and societal contexts ^11,12^. While higher SES has been held to be neuroprotective ^1–3^, ample evidence also exists for it being neuroselective ^13,14^. Both genes and environments vary with SES ^15^, and any observed relationship does not need to be causal in nature. Differences in SES-brain-cognition associations across cohorts have implications for whether relationships can be assumed to arise from direct or indirect effects of SES in early development or aging. More generally, different relationships across cohorts have implications for whether, when and how brain and cognitive function can be impacted by SES, or vice versa.

Here we address this question by investigating how SES variables in different cohorts originating in seven European countries, as well as in the US, relate to measures of brain structure and cognitive function. We test whether age-differences (child and adolescent development vs. adulthood/aging) and differences in sample origin (within Europe and Europe vs. US) are of importance to the relationships. A primary question is to what extent SES may exert influence on cognition through effects on brain structure through the lifespan, e.g. either affecting brain development or aging. It should be noted that neither causality nor direction of causality is given.

For instance, cognitive function could affect SES directly. People with higher cognitive ability may seek to have more education or income, and this may or may not lead to, or originate in, health behaviors that relate to brain volumes. Regardless of a possible bidirectional and complex nature, a relationship between SES and cognition may be mediated by brain characteristics. Thus, we tested to what extent brain variables could explain SES-cognition relationships.

The neural substrate for GCA is distributed across the brain ^8,16^. Also anatomically widespread associations between SES and neuroanatomical features have been reported ^5,17^. Hence, gross gray matter (GM) volume seems a good proxy for the brain foundations of SES-GCA relationships. GM volume is known to increase sharply along with cortical surface expansion in early childhood ^18^, and decrease in aging along with cortical thinning and subcortical volume reductions ^19^. Change in intracranial volume (ICV), on the other hand, comes to a halt after an initial period of development, and little if any age differences are seen after childhood ^20,21^. ICV therefore may serve as a proxy for maximal brain size ^22^. Hence, if SES variables are linked to ICV, this may be seen as a relationship intrinsic to neurodevelopment. If however, SES is related to GM volume in adult and aging populations when ICV is controlled for, then this may relate to variance in brain maintenance ^23^ or neurodegeneration.

As for sample origin, one debate has centered on possibly greater effects of variation in SES in US than in Europe ^11,24–26^. This could be the case if the extent of stratification by SES differs between US and Europe, or if SES variation is greater in the US ^11^. Different effects of SES could also to a greater extent reflect differences in opportunity for optimal development or maintenance of brain structure and cognition in US than in Europe. For instance, differences in income could be more linked to health and education in the US where higher education and health services are not provided as part of a free or minimal-cost welfare system in contrast to some European countries. It should be noted, however, that variation in socioeconomic inequalities, educational systems, and welfare states is also substantial across birth cohorts and within Europe ^27^. For instance, the UK provides a national health system, but while population health is worse in the US than in England, similar inequality in health by income have been found ^28^. Such income gradients may also apply to neurocognitive characteristics. Furthermore, the currently included cohorts are bound to vary in population representativeness, so while analyses here will illuminate differences across the specific cohorts studied, they may not readily be generalized to national differences more broadly.

We study multiple samples within the Lifebrain consortium ^29^, and also across other European and US databases, namely the UK Biobank (UKB) ^30,31^, the Human Connectome Project (HCP) ^32^, and the Adolescent Brain Cognitive Development (ABCD) study ^33,34^. We calculated per-site and across-site effect sizes for SES-brain-cognition relationships. The major goal of the Lifebrain consortium is to ensure a fuller exploitation, harmonization and enrichment of some of the largest longitudinal studies of age differences in brain and cognition in Europe. Hence a stream-lined analysis of possible differences in SES-brain-cognition relationships in these data sets, in combination with other European and US databases, will serve as an assessment of the effect sizes of these relationships, and how they differ across cohorts. Such an encompassing multi-national meta-analysis on SES-brain-cognition relationships across the lifespan is a novel undertaking.

Based on theoretical perspectives and evidence reviewed above, we hypothesized that SES-brain-cognition relationships would be found both in development and in adulthood/ aging. We expected the relationships to vary in strength across cohorts, regardless of age. We also expected differences between US and European cohorts, but both regional differences and differences in sample characteristics within each subset may be greater than general differences between continents. Based on the variable nature of previously reported relationships, we hypothesized that SES-cognition relationships could partly be explained by differences in brain structure.

By this multi-cohort approach we show that there is substantial heterogeneity in SES-brain-cognition relationships. This clearly demonstrates that SES does not exert influence on either brain or cognition, or vice versa, in any uniform way across cohorts. There were stronger positive relations between SES and brain structure in the US than in the European cohorts. ICV was more strongly related to SES than was GM volume controlled for ICV. This indicates a primarily developmental effect rather than neuroprotection in aging. Finally, ICV and GM volume explained part of the variance in both the education-GCA relationship and the income-GCA relationship.

## Results

### Regional cortical associations with GCA and SES

All statistical tests were two-sided. First, we validated that global GM volume is a better measure of brain structure in relation to GCA than regional volume, thickness or area. We ran general linear models (GLM) vertex-wise across the cortical surface, with GCA as predictor, and sex, study, age, age^2^ and ICV as covariates, for the participants for whom reconstructed surfaces were available (development < 20 years, n = 9,689; adulthood ≥ 20 years, n = 42,336, see SI for details). This analysis (Figure 1, see Supplementary Figure 1 for right hemisphere) showed extensive positive relationships across the cortical surface for volume and area. Bidirectional relationships were seen for thickness, especially in development, as expected due to ongoing developmental cortical thinning in this age-range. The volumetric results were most uniform in terms of direction of effects and a broad anatomical distribution. The same analyses were run using each SES variable as predictor, also showing widespread effects with most consistent results for volume (income, see Supplementary Figure 2; education, see Supplementary Figure 3). This suggests that global GM volume is a good summary measure of brain structure also in relation to GCA and SES. Further GM analyses were thus conducted on global GM volume, hereafter termed GM. Variation in GM and ICV in relation to age are shown in Supplementary Figure 4 across all cohorts.

**Figure 1.**
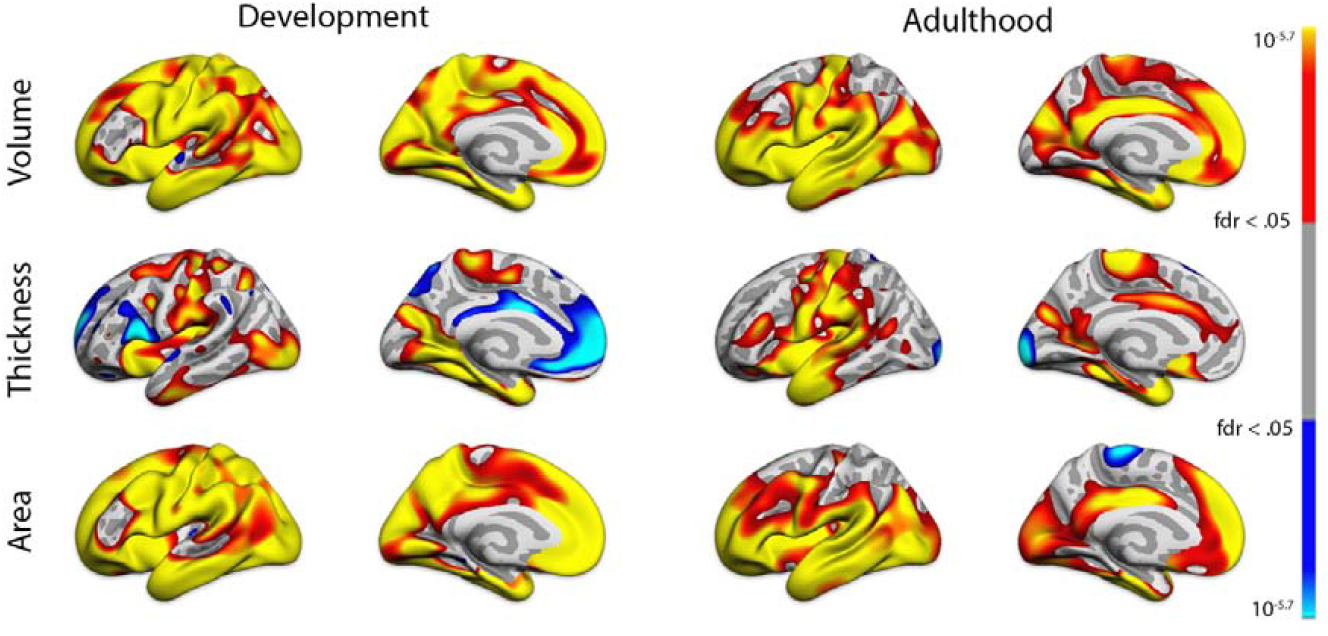
Effects of general cognitive ability (GCA) on cortical volume, area and thickness. Results corrected by false discovery rate < .05.

### Income and education

In all main analyses, variables of interest were adjusted for sex and age using a smoothing spline. To compare effect sizes across subsets of cohorts (development vs. adulthood, European vs. US), Wald tests for mean group differences between group-level meta-analytic estimates were run. We ran analyses on the cohorts grouped by age (development vs. adulthood) and in the full sample to obtain meta-analytic effect size estimates of both the grand-average relationships and the group-level relationships. For developmental cohorts, the parental education and income were used as SES.

As income data were given in various bins across studies, it is not possible to provide an informative measure of dispersion of income across studies. An overview of dispersion of years of education across studies is given in Supplementary Table 1. A Welch two group t-test showed that the variance in education was greater in European than US samples (t = 3.9567, df = 3.0835, p = 0.0274). The relationship between education and income was significantly positive overall (r = .30, 95% CI: .20 – .40), but heterogeneity was large (*Q*=2185.99, *p* < .0001, *I*^2^=99.29%, see Supplementary Figure 5). The association was in the positive direction in all 10 cohorts contributing both measures, and reached significance (as evidenced by CI not overlapping 0) for all but 1 cohort (NESDA). While apparently stronger relationships between income and education were found in US (r = .51, CI: .20 – .82) vs European (r = .25, CI: .18 – .31) cohorts, the difference was not significant (Z= - 1.611, p= 0.1100).

### Relationships of GCA with SES and brain structure

In analyses with total GM volume ICV was controlled for in addition to the other variables as listed above. Relationships of GCA to education, income, GM and ICV are shown grouped by developmental and adult cohorts in Figure 2. For plots grouped by European vs. US, see Supplementary Figure 6. The overall GCA-education correlation was r= .32 (CI: 0.22 – .41). The association was significantly positive in 9 of the 10 cohorts, as evidenced by CIs not overlapping 0. However, as the only two developmental cohorts included in this analysis showed very different effect sizes, the association was not significant across the developmental cohorts. The overall GCA-income correlation was r= .15 (CI: .06 – .24). The association was significantly positive in 6 of the 9 cohorts included, but was negative, although not significant, in 2 cohorts (BASE II, LCBC-dev), rendering the associations not significant in development. The overall GCA-GM correlation was r= .08 (CI: .04 – .11). The association was in the positive direction in all 11 cohorts included, but was significantly different from zero in only 4 (Whitehall, UKB, BASE II and ABCD). The overall GCA-ICV correlation was r= .15 (CI: .10 – .20). The association was in the positive direction in all 11 cohorts included, and was significantly different from zero for 8 (Whitehall, Betula and BASE II being the exceptions). Heterogeneity overall was large (GCA-education: Q=809.36, p <.001; GCA-income: I^2^=98.92%; Q=569.88, p < .0001; GCA-GM: I^2^=98.50%; Q=143.03, p < .0001; GCA-ICV: I^2^=86.00%, Q =52.95, p < .0001 I^2^=93.36%). In no case did the differences between developmental and adult cohorts in the GCA associations reach significance (all p >.25).

**Figure 2.**
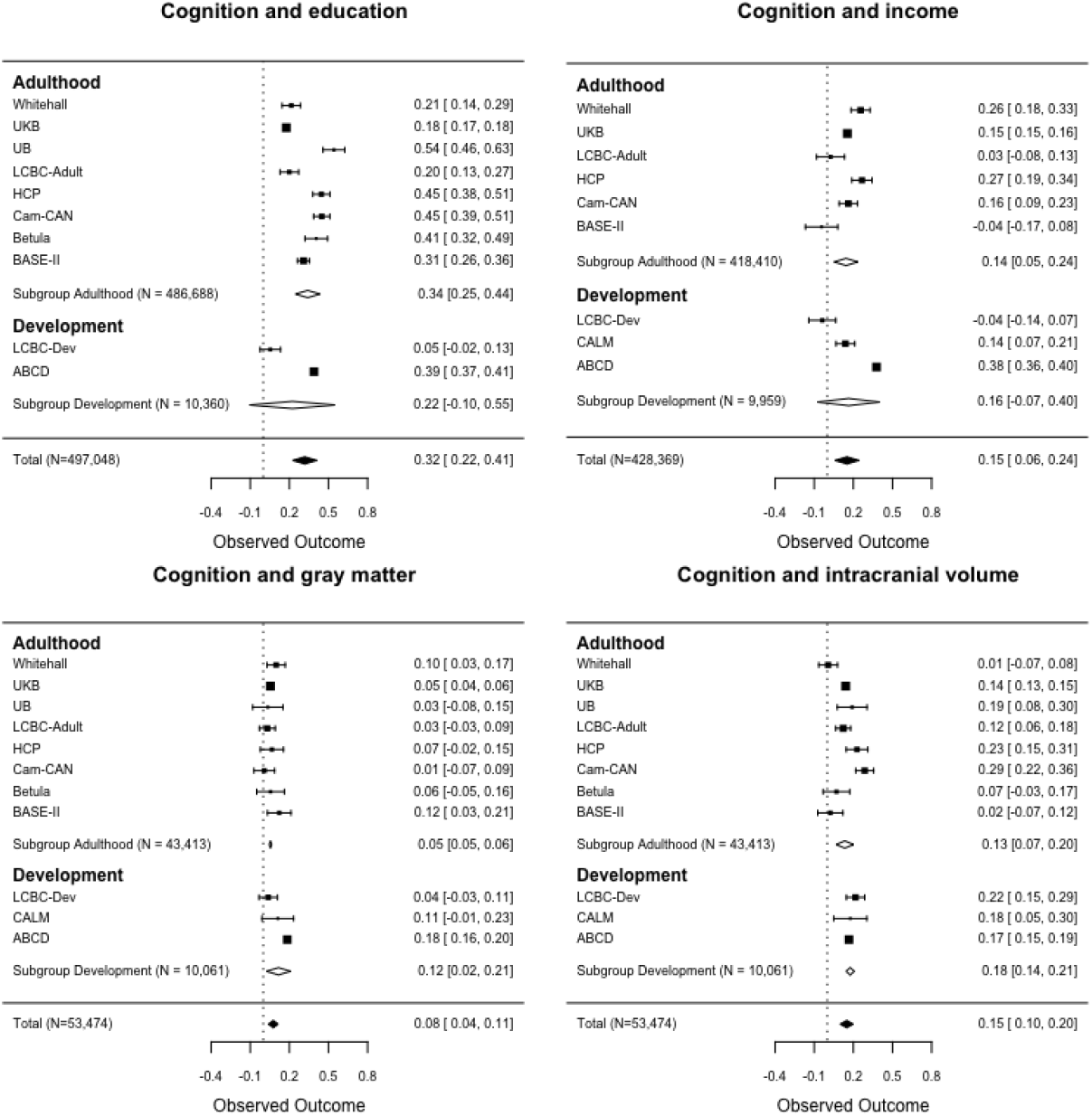
The associations of cognition (GCA) with education, income, GM and ICV. Forest plots show the individual observed effect sizes with corresponding 95% confidence intervals (CI). Diamonds represent the weighted average correlation estimate and its 95% CI with white diamonds representing the subgroup estimates and black diamonds the overall estimate. Numeric values of the cohort-specific and meta-analytic estimates are given in the right column. CI not spanning 0 indicates a significant (p<.05) relationship.

The difference in the GCA-education associations between European (r = .29, CI: .18 – .40) and US cohorts (r =.41 (CI: .36 – .46) was not significant (Z= –1.834, p= 0.067). However, the GCA-income association was significantly stronger in the US (r = .33, CI: .22 – .44) than in European (r = .10, CI: .02 – .18) cohorts (Z = −3.276, p= 0.0011). There was no significant difference in the GCA-GM associations adjusted for ICV between European (r = .05, CI: .05 – .06) and US (r =.13, CI: .02 – .25) cohorts (Z =-1.341, p = .18), or for the GCA-ICV associations (European: r = .14, CI: .08 – .20; US: r =.18, CI: .13 – .24; European-US difference: Z =-1.104, p = .27).

### Relationships between brain structure and SES

Relationships of GM and ICV with SES are shown grouped by developmental and adult cohorts in Figure 3. For plots grouped by European vs US, see Supplementary Figure 7. The overall GM-education correlation was r= 0.06 (CI: .01 – .11). The association was in the positive direction in 9 of the 12 cohorts included, but only significant in 3 (NESDA, HCP and ABCD). The effect was in the negative direction, although not significant, in two adult cohorts (Betula and Cam-CAN) and was numerically zero in an additional cohort (UB). There was no overall significant GM-income correlation (r= 0.05, CI: −.01 – .11). The association was in the positive direction in 6 of the 11 cohorts included, and significant in 5 (Whitehall, UKB, HCP, LCBC-Dev and ABCD), numerically zero in 1 (Cam-CAN), and in the negative direction in 4 cohorts (LCBC-adult, BASE II, HUBU, CALM). There was no significant difference in the GM-income association between developmental and adult cohorts (Z = −0.538, p = .5900). Heterogeneity was large for the GM-education (Q=248.19, p < .0001, I^2^=91.74%) and GM-income (Q= 251.14, p < .0001, I^2^=93.56%) associations.

**Figure 3.**
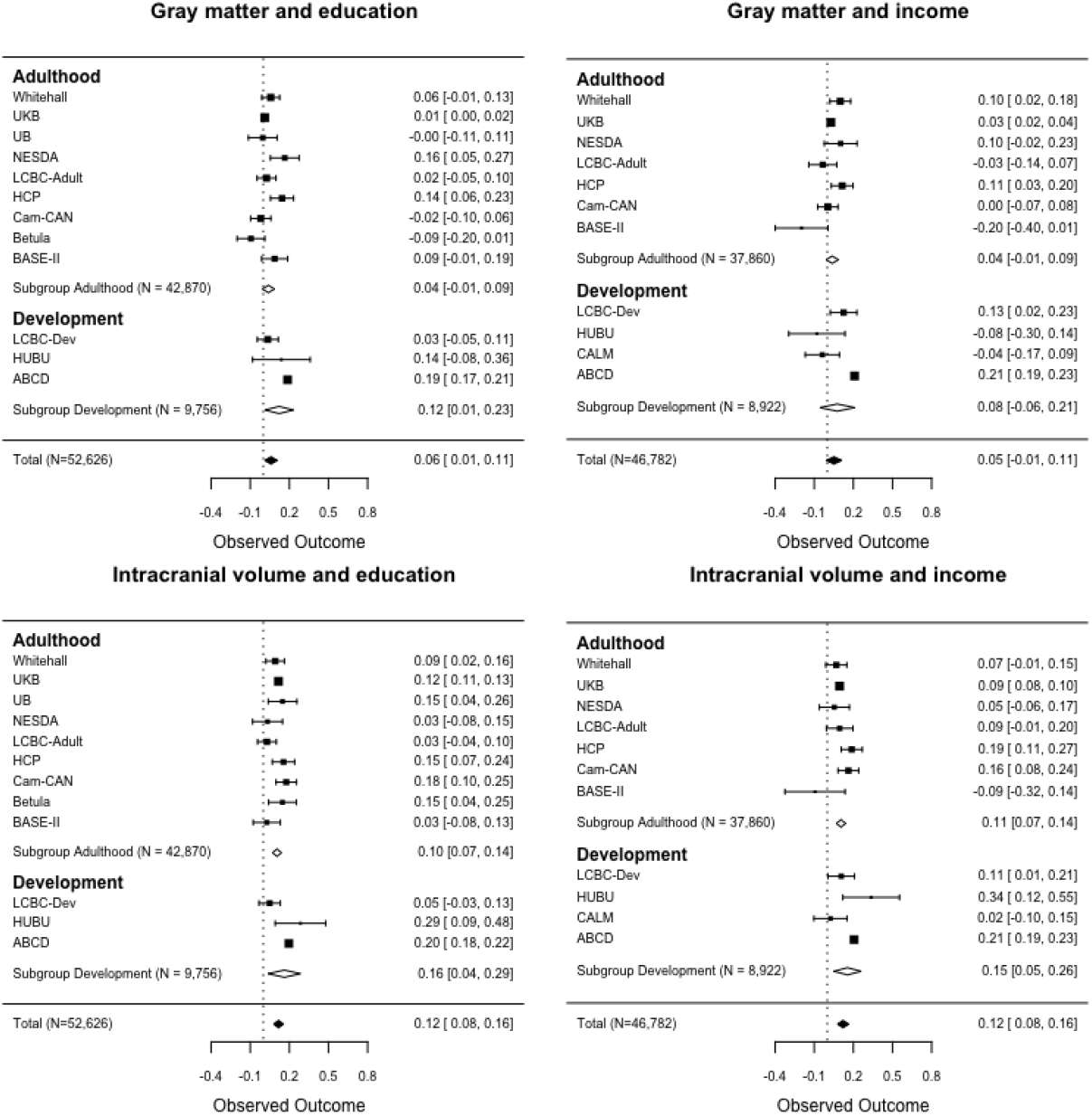
The associations of GM and ICV with education and income. Forest plots show the individual observed effect sizes with corresponding 95% CI. Diamonds represent the weighted average correlation estimate and its 95% CI with white diamonds representing the subgroup estimates and black diamonds the overall estimate. Numeric values of the cohort-specific and meta-analytic estimates are given in the right column.

Income was significantly more strongly associated with GM (Z= −2.763, p= .0057) in US (r = .17, CI: .08 – .26) than European cohorts (r = .03, CI: −.02 – .07). Education was also significantly more positively related to GM (Z= −7.987, p< 0.0001) in US (r = .18, CI: .16 – .20) than European cohorts (r = .03, CI: −.01 – .06).

The overall ICV-education correlation was r= .12 (CI: .08 – .16). All associations were positive, and significant relationships were observed in 8 of the 12 cohorts included (not in NESDA, LCBC-dev, LCBC-adult and BASE II). Heterogeneity was large (*Q*=72.44, *p* < .0001, P=85.66%). The ICV-education association was not significantly different (Z=-.858, p>.39) between developmental and adult cohorts, but was significantly greater (Z= −4.528, p< .0001) in US (r =.19, CI: .17 – .21) than European (r = .10, CI: .06, .14) cohorts. The overall ICV-income correlation was r= .12 (CI: .08 – .16). The associations were positive in 10 of 11 cohorts, being significant in 6 cohorts (UKB, HCP, Cam-CAN, LCBC-dev, HUBU and ABCD). Heterogeneity was large (Q=72.44, *p* < .0001, P=85.66%). The ICV-income association was not significantly different between developmental and adult cohorts (Z = – 0.876, p = .3800), but was significantly greater (Z= −9.583, p < .0001) in US (r =.20, CI: .18 – .22) than in European (r = .09, CI: .08 – .10) cohorts.

### Controlling for ethnicity and genetic ancestry

We addressed whether ethnicity or genetic ancestry factors (GAF) affected the relationships. It should be noted in this regard that there is substantial variation across cohorts in ethnicity, and the extent to which ethnic variance is present in the populations from which they were recruited also varies substantially. Controlling for either reported ethnicity (available for UKB, HCP, and ABCD, see Supplementary Figure 8 and 9) or GAF (available for UKB, LCBC-adult and LCBC-Dev, see Supplementary Figure 10 and 11) did not change the results substantially in the adult cohorts. In general, also relatively little change was observed in the relationships in the Norwegian developmental cohort (LCBC-Dev) when controlling for GAF, but the ICV-income relationship was no longer significant (GAF-adjusted r = .04, CI −.08 – .15; not GAF-adjusted r = .11, CI: .01 – .21). Relationships in the ABCD cohort appeared attenuated overall, albeit still significant when controlling for reported ethnicity. GCA-education and GCA-income relationships in ABCD appeared stronger when not controlling for GAFs (going from r =.39, CI: .37 – .41 to r = .28, CI: =.26 – .29 for GCA-education and from r = .38, CI: .36 – .40 to r = .24, CI: .22 – .26 for GCA-income). Associations between GCA and brain measures with SES variables in ABCD were the proportionally most attenuated overall after controlling for ethnicity (going from being in the range of r = +/-.20 to r =+/- . 10), but were still significant.

### The role of ICV

Differences in SES-ICV relationships and SES-GM relationships when ICV is controlled for have implications for the extent to which effects on the brain may be established in development or adulthood/aging. We found that education related more strongly (p = .044) to ICV (r = .12, CI: .08 – .16) than to GM controlled for ICV (r =.06, CI: .01 – .11). The same was the case for income (p = .0270) (with ICV: r = .12, CI: .08 – .16; with GM controlled for ICV: r = .05, CI: −.01 – 11). Notably, GCA was significantly more positively related to ICV than to GM controlled for ICV (p =.0294). GCA was also more positively related to education than to income (p = .0079). For details on these comparisons, see Supplementary Information (SI), Supplementary Figure 12.

### Effects of GM and ICV on SES-GCA relationships

We tested the extent to which the brain variables could explain part of the relations between cognitive ability and SES. We did this by testing the difference between the GCA-SES correlations adjusted for age and sex and the same correlations additionally adjusted for ICV, GM, or both. Overall, the correlations between GCA and SES were larger when not being adjusted for brain variables, especially for ICV. The GCA-education relationship was significantly more positive when not adjusting for ICV (p = .0034), GM (p = .0115) or ICV and GM combined (p = .0180). The GCA-income relationship was also significantly more positive across cohorts when not being adjusted for these brain variables (not adjusting. vs adjusting; for ICV: p = .0065; GM: p = .0428; ICV and GM: p = .0194). There was considerable variance across cohorts in the extent to which these variables altered the correlations. Only in the ABCD, HCP and the UKB cohorts, however, were SES-GCA associations significantly lower when controlling for any brain variable (see Supplementary Figure 13 for details).

## Discussion

This study shows substantial heterogeneity in SES-brain and cognition relationships across US and European cohorts encompassing all ages of the human lifespan. First, these results nuance the role of income in brain and cognitive development and aging in cohorts in industrialized countries, as uniformly positive effects were not the rule. As samples are highly heterogeneous and may have varying degrees of representativeness of the populations of origin, caution is warranted in interpretation of effects. Second, while education was as expected related to cognitive ability, and also showed some relationship to GM volume, a stronger relationship was observed for education and ICV. This may imply that associations between education and brain characteristics are grounded in neurodevelopment, as ICV changes relatively little after school-age is reached, and is known to stabilize between 10 years of age ^21^ and mid-adolescence ^20^. GM volume, on the other hand, for which less effect of education, and no overall effect of income, was found, shows substantial age differences across the lifespan, especially in older age ^35–37^. As years of education typically accumulate after ICV no longer increases, a direct effect of education on ICV in adulthood is improbable. This is also consistent with the association between ICV and education being significantly greater in developmental than in adult cohorts. Having more educated parents – or a correlate thereof, could have a facilitating effect on brain development in childhood and possibly adolescence. While individual education often has been seen as boosting development and being neuroprotective ^1–3^, ample evidence also exists for it being neuroselective ^13,14^. The development-neuroprotection account implies a causal effect of socioeconomic status, whereas in a strict neuroselective view, education and income would rather be markers or proxies of some other favorable, putatively genetic, trait ^13,14^. A higher ICV could in part reflect causal factors in driving years of education in adulthood. However, one needs to keep in mind that ICV has shown very high heritability, up to .88 in some studies ^38^, and genetic pleiotropy of ICV and education may be likely.

As this study was performed on cross-sectional data, conclusions regarding change in brain and cognition cannot be drawn. Less knowledge exists on SES-brain structure relations in midlife and aging, but a relatively large US study found that community disadvantage in midlife was associated with reduced cortical tissue volume, cortical surface area, and cortical thickness, but not subcortical morphology ^39^. Hence, while most focus has been on development, there is no reason to believe that overall relations between SES and brain structure is confined to young samples. Indirect evidence for this also comes from epidemiological data, where lower SES is associated with greater risk of dementing diseases characterized by brain atrophy or lesser neuroanatomical volumes ^1^. However, the fact that SES-brain-cognition relationships are found in aging cohorts, should not be taken to indicate that they operate in aging specifically, rather than in a stable manner, perhaps as an intercept effect across the lifespan. The current results do not support a neuroprotective account, where higher SES serves to mitigate cognitive decline or GM atrophy in aging. Neither education nor income were consistently positively associated with ICV-adjusted GM volumes, and relationships with GM and cognitive ability did not significantly differ in developmental and aging cohorts.

### Comparison between European and US samples

The question of whether SES-brain-cognition relationships differ across cohorts and societies has also been highlighted by other types of studies. Evidence for an SES-genotype interaction on cognitive ability has been found, in terms of suppression of heritability with lower SES ^11,24^. Recently, such effects and as their possible current absence in European and presence in US samples have been debated ^11,25,26^. Our results show substantial heterogeneity of SES-brain-cognition relationships across cohorts also within Europe, and even from the same country, as can be appreciated by differing effect sizes across UK cohorts.

However, there were significantly different effects of income on cognition, and of income and education on GM and ICV between US and European samples. These differences all point to stronger positive relationships between SES and brain and cognition in the US than in the European samples. Large US studies on developmental samples, one of which included here, have shown broadly distributed associations between SES and brain structure ^5,17^. One recent European longitudinal study found widespread associations between a composite SES measure and cortical surface area at age 14, with independent contributions from polygenetic scores for education ^40^. Some US studies have found the strongest associations, with especially lower regional neuroanatomical volumes, in children living in poverty ^41,42^. Somewhat less evidence is available from European cohorts, although such associations have also been found in young cohorts in Germany and France with large variation in SES ^43,44^. However, in a Norwegian sample ^8^, including a subset of the one entered in present analyses, no associations were found between income or education and regional cortical area. The current finding of US-European differences is thus not completely unexpected. However, it should be emphasized that the currently included cohorts will vary in representativeness of the populations from which they were drawn. For both US cohorts, efforts were made to recruit participants reflecting the ethnic and sociodemographic composition of the population ^34,45^. While this was also the case for many of the European cohorts, see e.g. ^31,46,47^, the differences across US and European cohorts may still to some extent reflect more diverse sociodemographic backgrounds in the US than European studies. However, it should be noted that in terms of years of education, the variance was greater in European than US samples.

As for SES and GCA, a few meta-analyses exist ^7,48,49^, all reporting relationships between intelligence quotient (IQ), income, and education. In the most comprehensive meta-analysis so far, differences in IQ-SES relationships in the U.S. vs. other Western societies were not supported ^7^. This is in line with our findings for education, but we note that a stronger positive relationship between income and cognitive ability was observed in US samples. Income was not consistently related to cognitive ability across the present cohorts. The most positive relationships were found in the US and UK cohorts, while there were other European cohorts in which no relationships were seen. Hence, the results highlight that income may not be related to cognitive function in a global way.

### The role of brain structure in SES-cognition relationships

Finally, the current results did not only yield support for SES-cognition and SES-brain relationships, but also showed that variance in brain structure, i.e. ICV and GM independently of ICV, explained part of the education-cognition and income-cognition relationships across cohorts. This was indicated by the fact that adjusting for either brain metric significantly weakened the relationships.

### Limitations and future directions

The current study has a number of limitations. Other relations could be uncovered with less general metrics than education, income, GM and ICV. For instance, occupation, subjectively perceived social rank, cortical thickness and area could be more refined measures. Still, the vertex analyses showed that both for GCA and SES, effects were anatomically widespread and more consistently related to volume than thickness or area, suggesting that GM volume is a sensitive measure of brain structure for our purpose. Further, ethnicity or GAFs were not included as covariates in all analyses. While some of the cohorts have no or minor ethnic variation, others have more (see SI). Analyses in select big cohorts did overall indicate, however, that the relationships in most cases remained significant when controlling for ethnicity or GAFs. Furthermore, the measures for some of the constructs studied here are quite heterogeneous. For instance, estimates of GCA were obtained with different tests of crystallized and fluid ability. The comparison of European and US cohorts is limited by the fact that we have only two US samples. In part, the same limitation goes for the relatively few developmental relative to adult and aging samples. As cohorts are not invariably representative of the societies from which they are recruited, further interpretation of these possible differences is not warranted here. However, the substantial heterogeneity should prompt researchers to carefully examine relationships before SES indicators are used as covariates of no interest. Such practices may otherwise suppress or inflate variance in relationships of interest in unpredictable ways.

### Conclusions

There is substantial heterogeneity in the relationships of SES to brain and cognition across major European and US cohorts. Based on these results, it is not likely that the effects of SES on cognition are grounded in neuroprotective effects on GM volume in aging. Rather, SES relations established in brain development may be seen through the lifespan. In this regard stronger relationships of income and education to neuroanatomical volumes were found in US than European cohorts, pointing to SES not signifying the same across different populations from industrialized countries. The present results also indicate that part of the relations between SES and cognition may be mediated by SES-brain relationships. The findings have implications for our understanding of whether, when and how SES may impact brain and cognition.

## Online Methods

### Samples

All samples were recruited to be community-dwelling participants, some were convenience samples, whereas others were contacted on the basis of populations registry information. While we do not believe development ends at a particular point, for simplicity we here use the terms “development(al)” for the child and adolescent cohorts and “adult(hood)” for the cohorts with participants 20 years of age and above. Demographics of the samples are given in Table 1, see SI for details. For a visual representation of the age-distributions of the samples, see Supplementary Figure 14.

**Table 1.**
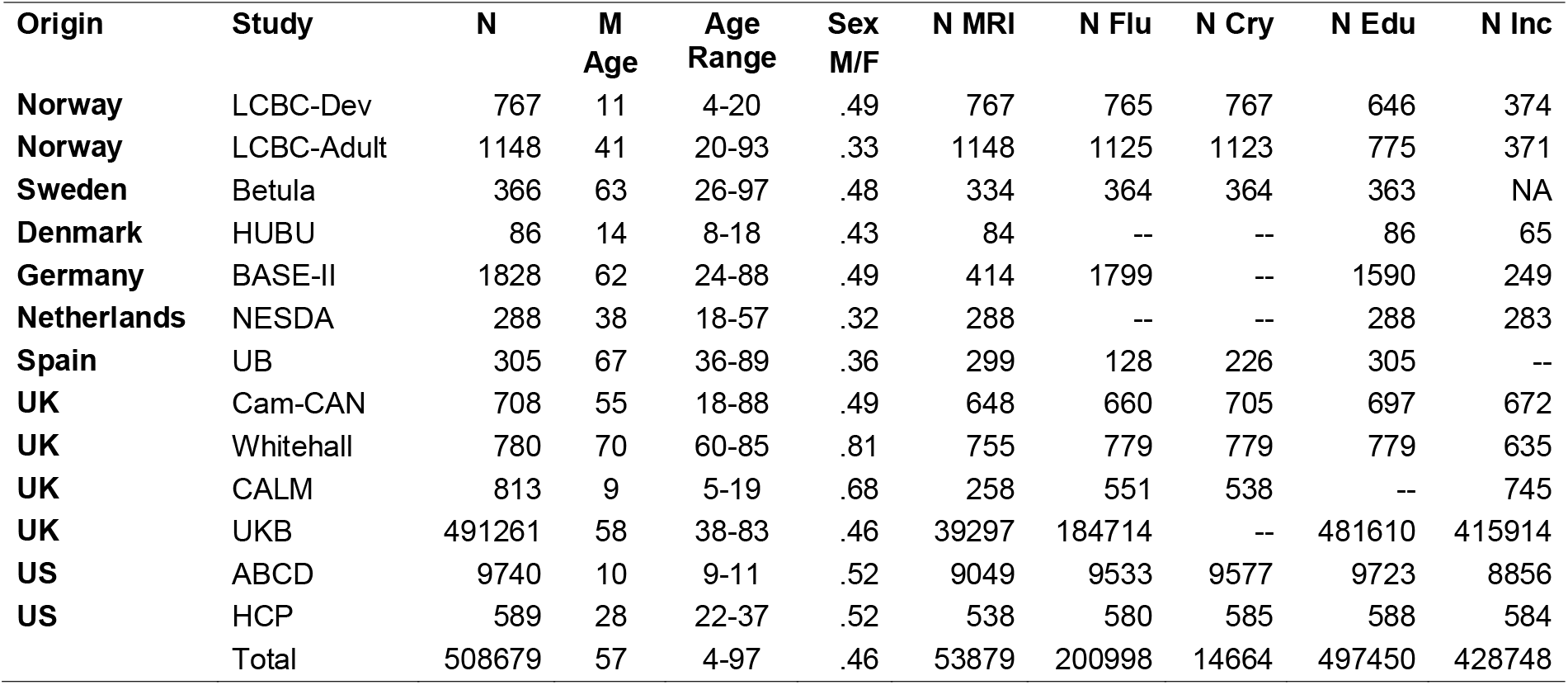
Overview of sample characteristics of included cohorts. Fluid = measures of fluid cognitive ability, Cry = Measures of crystallized cognitive ability, Edu = measures of education, Inc = measures of income. Sex M/F refers to sex ratio, male proportion.

### Lifebrain subsamples

The samples were derived from the European Lifebrain project (http://www.lifebrain.uio.no/) ^29^, including participants from major European brain studies: Berlin Study of Aging II (BASE II) ^46,50^, the BETULA project (^47^, the Centre for Attention, Learning and Memory study (CALM) ^51,52^, the Cambridge Centre for Ageing and Neuroscience study (Cam-CAN) ^53^, the Brain maturation in children and adolescents study (HUBU) ^54^, Center for Lifebrain Changes in Brain and Cognition longitudinal studies (LCBC) ^8,55^, the Netherlands Study of Depression and Anxiety (NESDA) ^56^, the University of Barcelona brain studies (UB) ^57–59^ and the Whitehall II Imaging sub-study (WH II Imaging) ^60^. In total, data from 7,089 participants (5-96 years of age) were included from the Lifebrain cohorts. However, all participants and all cohorts did not contribute all categories of data, as detailed in Table 1 and SI. Importantly, MRI-derived ICV and GM measures were available for 4,995 participants from the Lifebrain cohorts.

#### UKB

The UK Biobank (UKB) recruited 502,649 participants aged 37 to 73 years from 2006 to 2010 ^61^. Ethical approval was obtained from the National Health Service National Research Ethics Service (Ref 11/NW/0382) and all participants provided written informed consent. Here, the dataset released February 2020 was used, consisting of 502,507 participants, of whom 40,682 had undergone MRI scanning. After applying exclusion criteria (see SI), 491,261 had sufficient valid information to be included in the final analyses, of whom 39,297 had a valid MRI.

#### HCP

The Human Connectome Project (HCP) is funded by the US National Institute of Health (NIH) (http://www.neuroscienceblueprint.nih.gov/connectome/). The consortium led by Washington University and the University of Minnesota (the ‘WU-Minn HCP Consortium’) aims to study brain connectivity and function with a genetically informative design in 1,200 individuals using four MR-based modalities plus MEG and EEG. Behavioral and genetic data are also acquired from these participants. After application of exclusion criteria, 538 participants with MRI were included. For further information, see SI.

#### ABCD

The Adolescent Brain Cognitive Development (ABCD) study aims to track human brain development from childhood through adolescence ^33^. ABCD has recruited > 10,000 9-10 years olds across 21 US sites with harmonized measures and procedures, including imaging acquisition https://abcdstudy.org/scientists-workgroups.html and processing 62. A goal of the ABCD study is that its sample should reflect, as best as possible, the sociodemographic variation of the US population ^34^. For ABCD, the dataset release 2.0.1 was used, consisting of 11,875 participants at baseline, of whom 9,740 had sufficient valid information to be included in the final analyses (9,049 with MRI).

### General procedures

We used all available Lifebrain cohorts that provided at least two of the constructs of interest: GCA (crystallized and/or fluid intelligence), SES (income and/or education), and brain structure (GM volume and ICV), resulting in a total of 10 Lifebrain studies. Of the Lifebrain studies, 9 provided measures of education, 8 of income, 9 of brain structure, and 8 of crystallized and/or fluid intelligence, used to compute measures of GCA. In addition, analyses were performed on UKB, HCP, and ABCD.

For each study, we gathered all cognitive tests that we considered measuring fluid and/or crystallized intelligence. There were multiple tests for GCA in each cohort. Using principal component analysis, we reduced GCA to its first principal component. With this approach, we could analyze the correlations of four constructs of interest that we refer to as GCA, income, education and neuroanatomical volume. For details on how these constructs were recorded per study, see SI. For measures of neuroanatomical volumes, we gathered FreeSurfer-based estimates of total GM volume and ICV. We meta-analyzed Spearman rank-order correlations with bootstrapped standard errors based on 1,000 replications each. The bootstrapped standard errors served as weights for the meta-analysis. For GM and ICV, we ran separate regressions for each cohort predicting volume by age and checking whether absolute residuals exceeded a relatively liberal four standard-deviations criterion. If so, the respective participants were entirely deleted from the following analysis. For details, see SI.

### Magnetic resonance imaging acquisition and analysis

T1 weighted structural scans were acquired at Siemens (Erlangen, Germany), Philips and GE scanners at the various sites. Further information on MRI scanning and processing is given in SI. Images were processed with FreeSurfer, mainly version 6.0 (https://surfer.nmr.mgh.harvard.edu/) (FreeSurfer 5.3 was used for Whitehall II, HCP and ABCD). Because FreeSurfer is almost fully automated, to avoid introducing possible site-specific biases, gross quality control measures were imposed and no manual editing was done.

## Statistics

Meta-analyses were computed based on the primary outcome of a single effect size *r*, the pair-wise Spearman correlations among constructs. We used pairwise complete observations to compute correlations. When constructs had more than one indicator, we used principal component analysis as dimensionality reduction techniques to obtain factor score estimates. In order to obtain PCA estimates from missing data, missing data have to be imputed. Missing data in GCA were imputed using the regularized iterative PCA algorithm (with a single component) as implemented in the R package missMDA ^63^. Meta-analytic estimates of correlations and their precisions were obtained from the metafor package ^64^. As our primary outcome of interest is the latent correlation of pairs of constructs of interest (crystallized/fluid intelligence, income, education, and neuroanatomical volumes), effect size estimates were weighted by their inverse bootstrapped standard error (which implicitly considers sample size differences among cohorts). We report mean effect size and 95% CI for each study and for the meta-analytic effect size estimates. If the 95% CI did not include zero, the null hypothesis of no correlation could be rejected at a *α* level of .05. The meta-analysis was based on a random-effects model ^65^, in which both the within-study variance and the between-study variance form the variance component used to calculate study-specific weights ^66^. In contrast to a fixed-effects model, it is often described as more conservative. Importantly, the randomeffects model accounts for between-study heterogeneity, *τ*^2^, which itself is an outcome of interest for our analysis. To describe and test the heterogeneity in our results, we report *I*^2^, the ratio of between-study heterogeneity, *τ*^2^ over observed variability ^67^. *I*^2^ can be considered a standardized effect size estimate of heterogeneity or inconsistency across studies with larger values meaning a presence of more heterogeneity. To assess whether observed heterogeneity in the estimated correlations across studies are compatible with chance alone, we also report *p* values of Cochrane’s *Q* test of heterogeneity.

### Demographic measures

For all samples, age was measured in years and months, and converted to a three-decimal numeric value for analyses. Sex was coded as 0 for males and 1 for females. For details on how education and income was recorded, see SI. In general, estimates of parental education and income were used for developmental samples, whereas participant income and education were recorded for adult samples.

### Cognitive tests

For GCA, national versions of a series of batteries and tests were used, see SI for details. These included tests from the Wechsler Abbreviated Scale of intelligence (^68^ LCBC, CALM), Wechsler Primary and Preschool Scale of Intelligence III (^69^ LCBC–below age 6.5 years), the Wechsler Adult Intelligence Scale R/ III/ IV (^70,71^ UB, Betula, Whitehall II), Wechsler Individual Achievement Test (^72^ CALM), Test of Premorbid Functioning (^73^ Whitehall II), Cattell Culture Fair (^74^ Cam-CAN), National Adult Reading Test (^75^ UB), NIH toolbox (^76^ ABCD, HCP), as well as local batteries.

## Acknowledgements and funding information

The Lifebrain project is funded by the EU Horizon 2020 Grant agreement number 732592 (Lifebrain). In addition, the different sub-studies are supported by different sources: LCBC: The European Research Council under grant agreements 283634, 725025 (to A.M.F.) and 313440 (to K.B.W.), as well as the Norwegian Research Council (to A.M.F., K.B.W.), The National Association for Public Health’s dementia research program, Norway (to A.M.F). Betula: a scholar grant from the Knut and Alice Wallenberg (KAW) foundation to L.N. Barcelona: Partially supported by Spanish Ministry of Science, Innovation and Universities (MICIU/FEDER; RTI2018-095181-B-C21) research grant, and also supported by an ICREA Academia 2019 grant award; by the California Walnut Commission, Sacramento, California. BASE-II has been supported by the German Federal Ministry of Education and Research under grant numbers 16SV5537/ 16SV5837/ 16SV5538/ 16SV5536K /01UW0808/ 01UW0706/ 01GL1716A/ 01GL1716B, the European Research Council under grant agreement 677804 (to S.K.). The infrastructure for the NESDA study (www.nesda.nl) is funded through the Geestkracht program of the Netherlands Organisation for Health Research and Development (ZonMw, grant number 10-000-1002) and financial contributions by participating universities and mental health care organizations (VU University Medical Center, GGZ inGeest, Leiden University Medical Center, Leiden University, GGZ Rivierduinen, University Medical Center Groningen, University of Groningen, Lentis, GGZ Friesland, GGZ Drenthe, Rob Giel Onderzoekscentrum). HUBU has been supported by the Danish council of Independent Research | Medical Sciences (grant numbers 09-060166, 0602-02099B) and the Lundbeck Foundation (grant number R32-A3161). The Cambridge Centre for Ageing and Neuroscience (Cam-CAN) was supported by a programme grant from the UK Biotechnology and Biological Sciences Research Council (grant number BB/H008217/1) and by continued intramural funding from the UK Medical Research Council to the Cognition & Brain Sciences Unit in Cambridge. Work on the Whitehall II Imaging Substudy was funded by the UK Medical Research Council (G1001354) and the HDH Wills 1965 Charitable Trust (Nr: 1117747) to K.P.E. The Wellcome Centre for Integrative Neuroimaging is supported by core funding from award 203139/Z/16/Z from the Wellcome Trust. Data were provided [in part] by the Human Connectome Project, WU-Minn Consortium (Principal Investigators: David Van Essen and Kamil Ugurbil; 1U54MH091657) funded by the 16 NIH Institutes and Centers that support the NIH Blueprint for Neuroscience Research; and by the McDonnell Center for Systems Neuroscience at Washington University. Data used in the preparation of this article were [in part] obtained from the Adolescent Brain Cognitive Development (ABCD) Study (https://abcdstudy.org), held in the NIMH Data Archive (NDA). This is a multisite, longitudinal study designed to recruit more than 10,000 children age 9-10 and follow them over 10 years into early adulthood. The ABCD Study is supported by the National Institutes of Health and additional federal partners under award numbers U01DA041022, U01DA041028, U01DA041048, U01DA041089, U01DA041106, U01DA041117, U01DA041120, U01DA041134, U01DA041148, U01DA041156, U01DA041174, U24DA041123, U24DA041147, U01DA041093, and U01DA041025. A full list of supporters is available at https://abcdstudy.org/federal-partners.html. A listing of participating sites and a complete listing of the study investigators can be found at https://abcdstudy.org/scientists/workgroups/. ABCD consortium investigators designed and implemented the study and/or provided data but did not necessarily participate in analysis or writing of this report. This manuscript reflects the views of the authors and may not reflect the opinions or views of the NIH or ABCD consortium investigators. The ABCD data repository grows and changes over time. The ABCD data used in this report came from doi:10.15154/1506087.

Part of the research was conducted using the UK Biobank resource under application number 32048.

## Author contributions

Kristine B. Walhovd, Anders M. Fjell, Ulman Lindenberger, Lars Nyberg, Paolo Ghisletta, Klaus P. Ebmeier, Christian A Drevon, David Bartrés-Faz, Andreas M. Brandmaier conceived the idea and designed the study.

Andreas M. Brandmaier performed the meta-analyses.

Øystein Sørensen, Inge Amlien, Anders M. Fjell and Kristine B. Walhovd contributed to the statistical analyses.

Kristine B. Walhovd drafted the manuscript.

Yunpeng Wang conducted the genetic analyses.

Athanasia M. Mowinckel, Inge K. Amlien, Kristine B. Walhovd, Anders M. Fjell, Sandra Düzel, William Baaré, Rogier A. Kievit, Richard N. Henson, Kathrine Skak Madsen, Cristina Solé-Padullés, Sara Pudas, Lars Nyberg, Enikő Zsoldos, Klaus P. Ebmeier, Brenda Penninx, Laura Nawijn contributed to the acquisition and/ or preparation the data.

All authors provided critical feedback and editing of the final manuscript.

