## Supplemental Information for "Education and income show heterogeneous relationships to lifespan brain and cognitive differences across European and US cohorts"

**Supplementary Information**

**Supplementary Results**

**Cortical vertex-wise analyses**

We ran vertex-wise analyses to assess regional variation in the relationships between cortical structure and the measures of interest - GCA and each of the SES variables (income and education) - and whether cortical area or thickness showed more consistent relationships than volume. Cortical surfaces were reconstructed from the same T1-weighted anatomical MRIs used for global GM volume by use of FreeSurfer v6.0 (<https://surfer.nmr.mgh.harvard.edu>/https://surfer.nmr.mgh.harvard.edu)^1-4^, yielding maps of cortical area, thickness and volume. Surfaces were smoothed with a Gaussian kernel of 15 mm full-width at half-maximum. Three general linear models were run, for each of the variables GCA, income and education in turn as predictors, with sex, study, age, age^2^ and ICV as covariates. For consistency across analyses, the results were thresholded at p <.01 (uncorrected). In most cases, this was a stricter threshold than when FDR-corrections (FDR < .05) were applied.

Analyses were run for only those participants where reconstructed surface data were available at the research server in Oslo. Thus, the vertex-results are from a smaller sample than the total sample used for the main analyses, limited to the samples LCBC-dev, BASE-II, Betula, Cam-CAN, LCBC-adult, UB, ABCD, HCP and UKB. The results are shown in Supplementary Figure 1, 2 and 3.

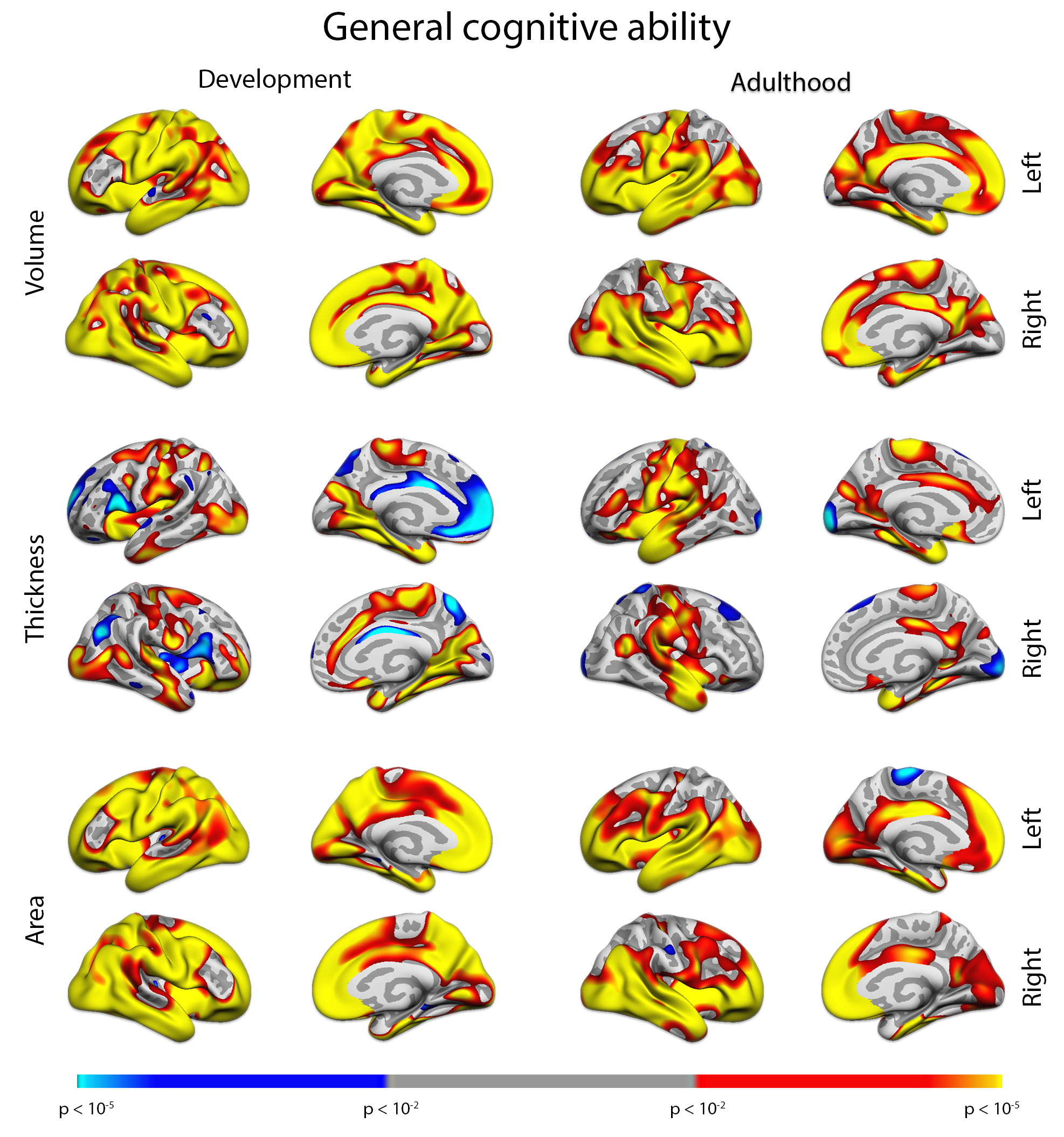

**Supplementary Figure 1**

Relationship between GCA and cortical morphometry. Development (n = 9,689) is shown to the left and adulthood (n = 42,339) to the right. From top to bottom rows, volume, thickness and area results are shown, respectively.

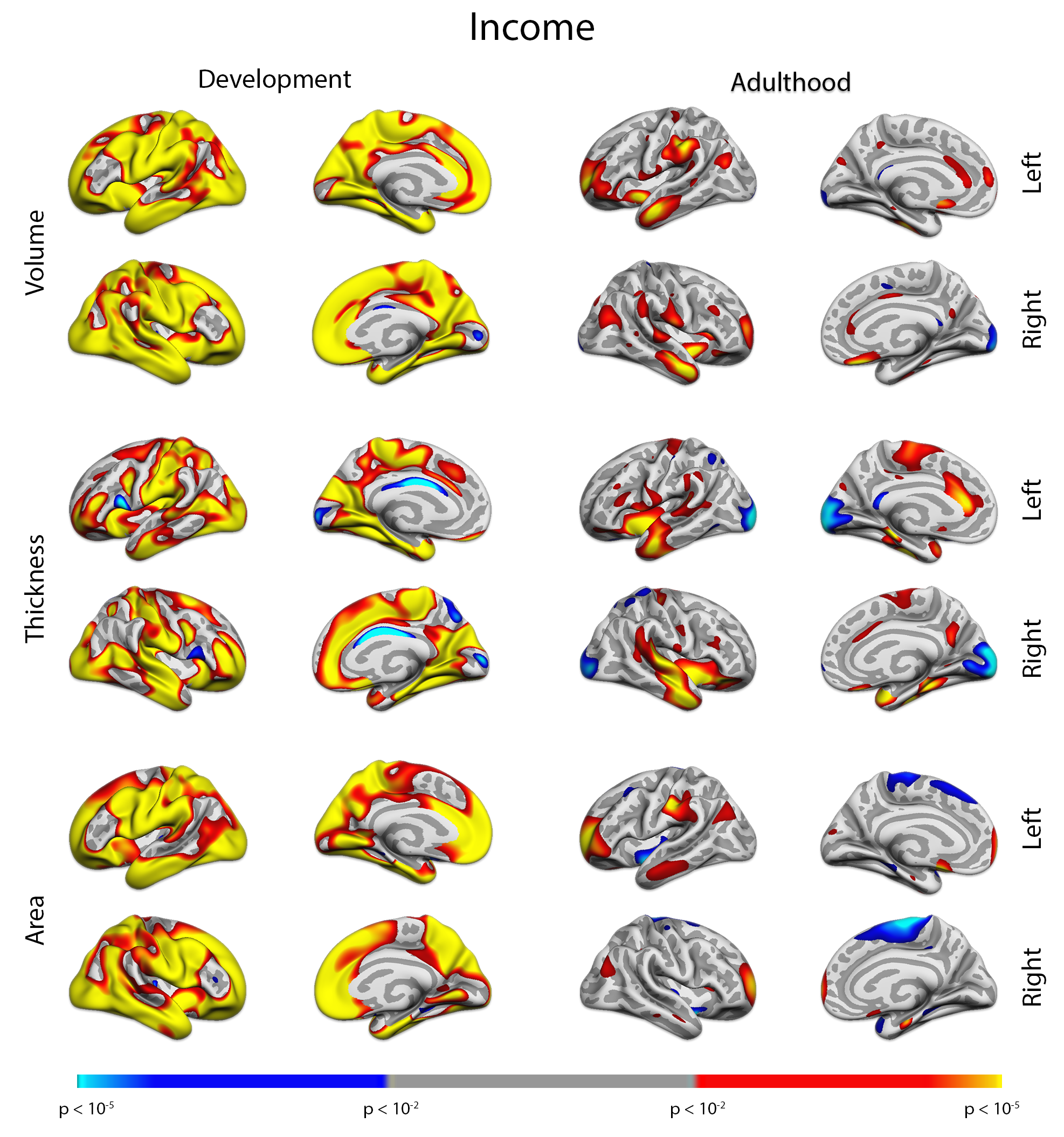

**Supplementary Figure 2**

Relationship between Income and cortical morphometry. Development (n = 8,505) is shown to the left and adulthood (n = 42,336) to the right. From top to bottom rows, volume, thickness and area results are shown, respectively.

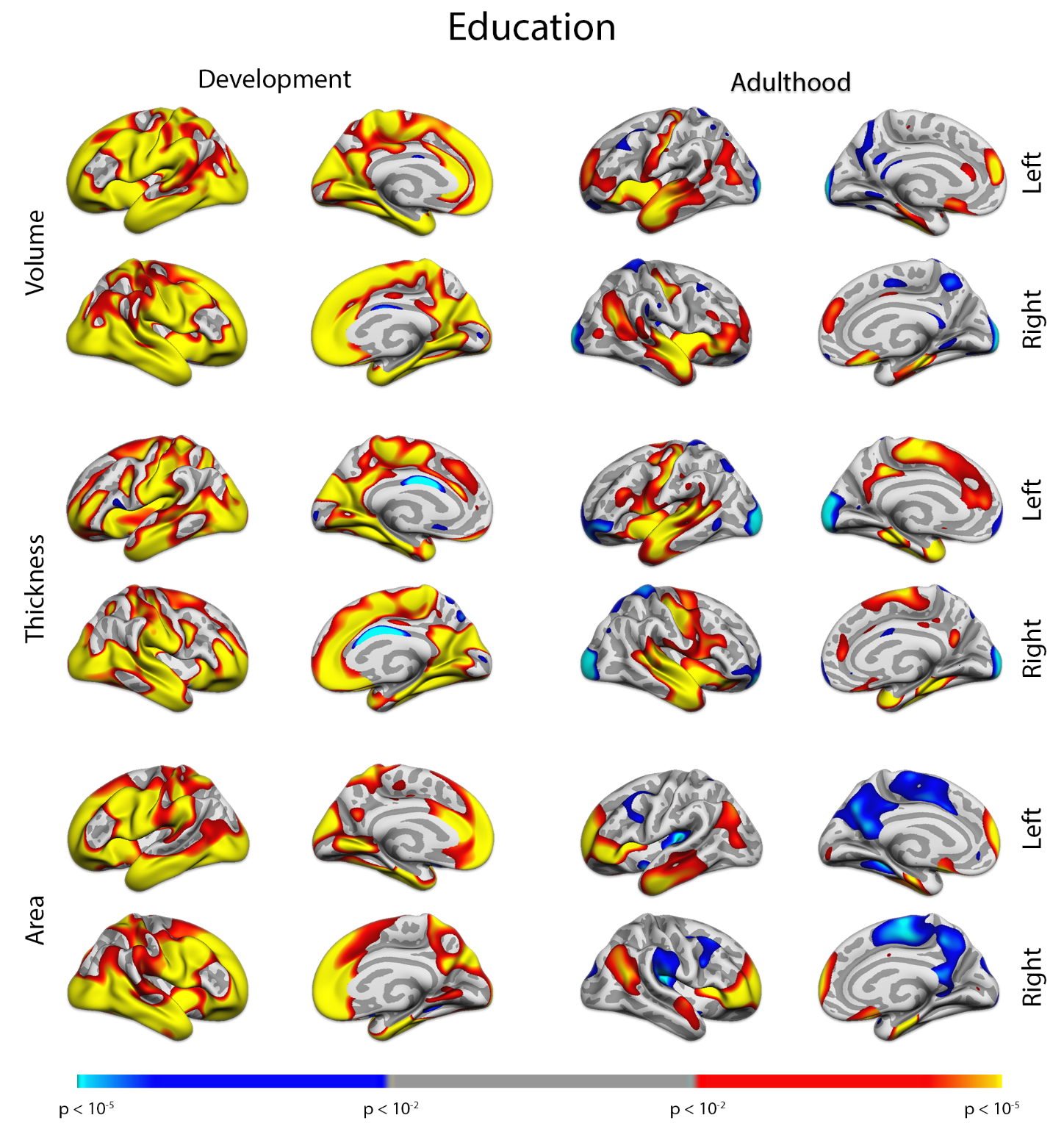

**Supplementary Figure 3**

Relationship between Education and cortical morphometry. Development (n = 9,550) is shown to the left and adulthood (n = 41,244) to the right. From top to bottom rows, volume, thickness and area results are shown, respectively.

| **GM** | **ICV** |  |
| --- | --- | --- |
| 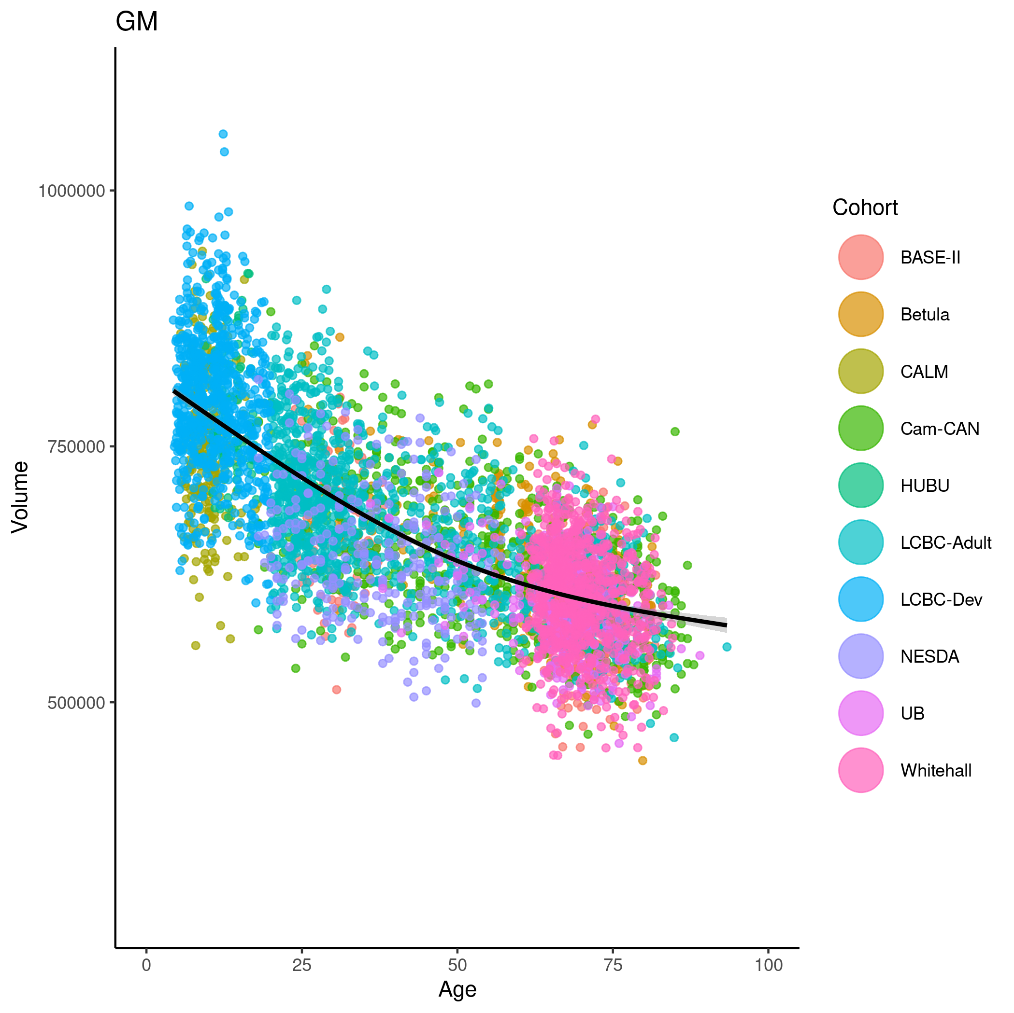 | 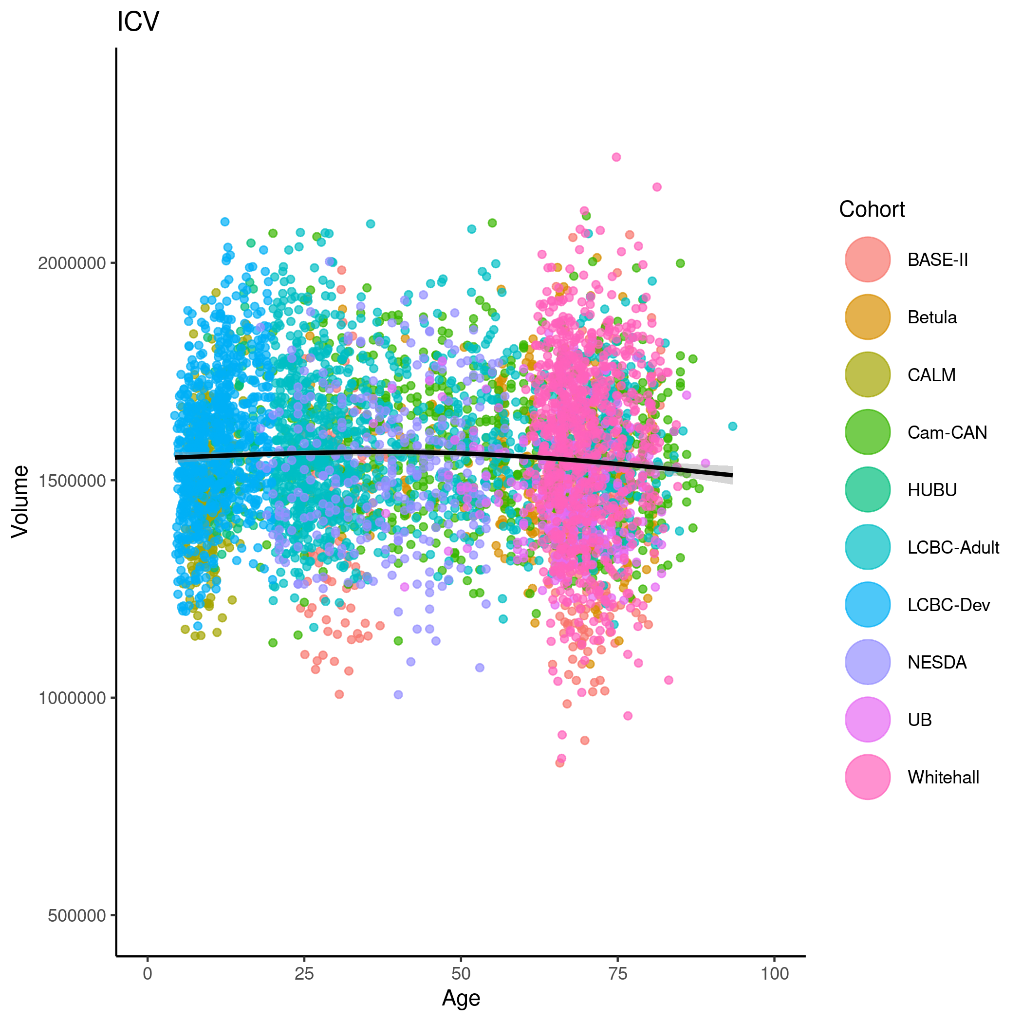 | 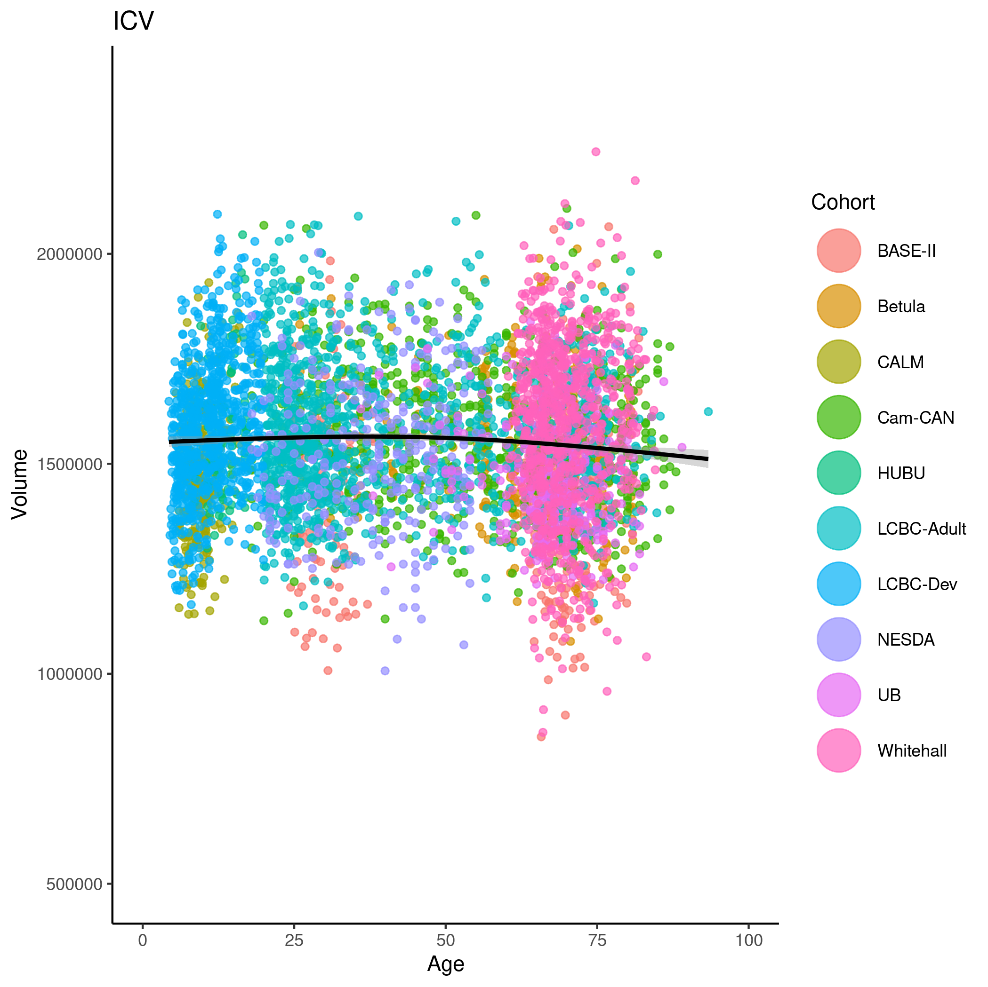 |
| 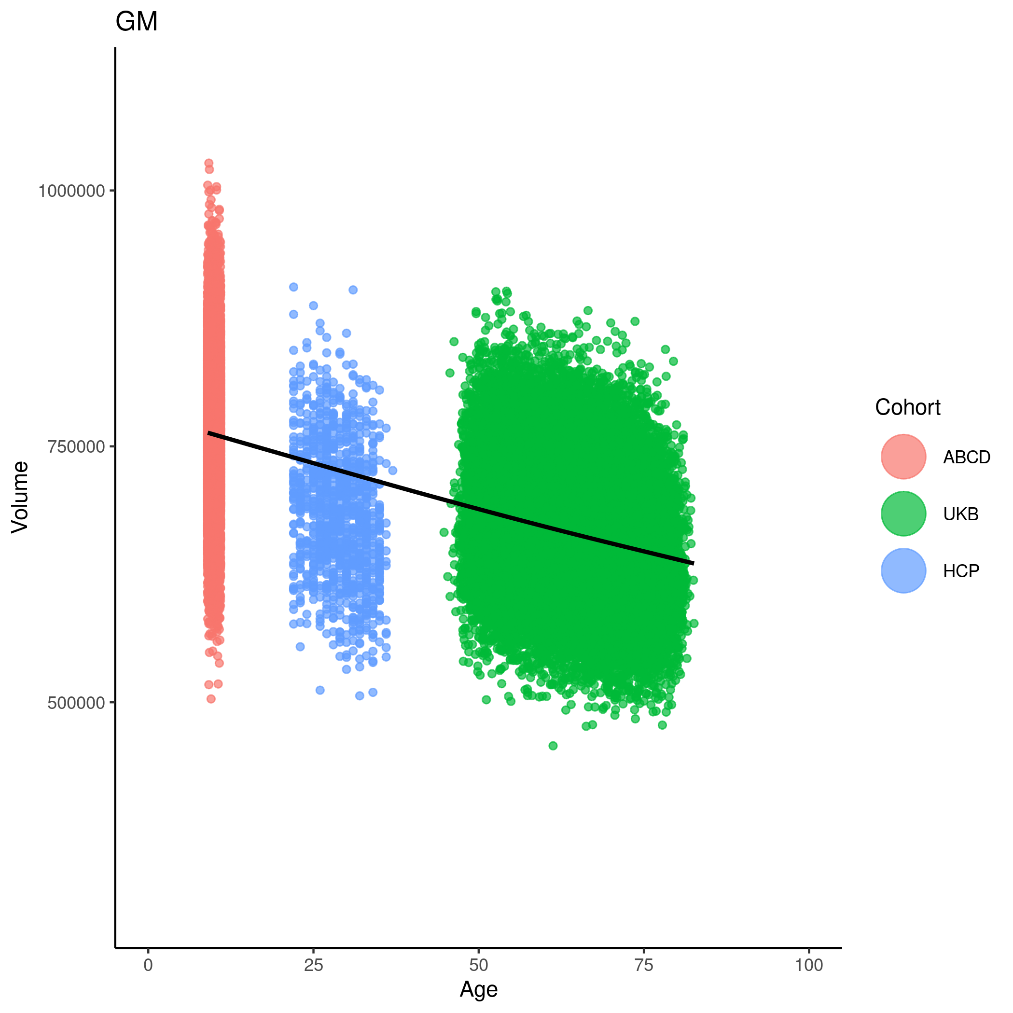 | 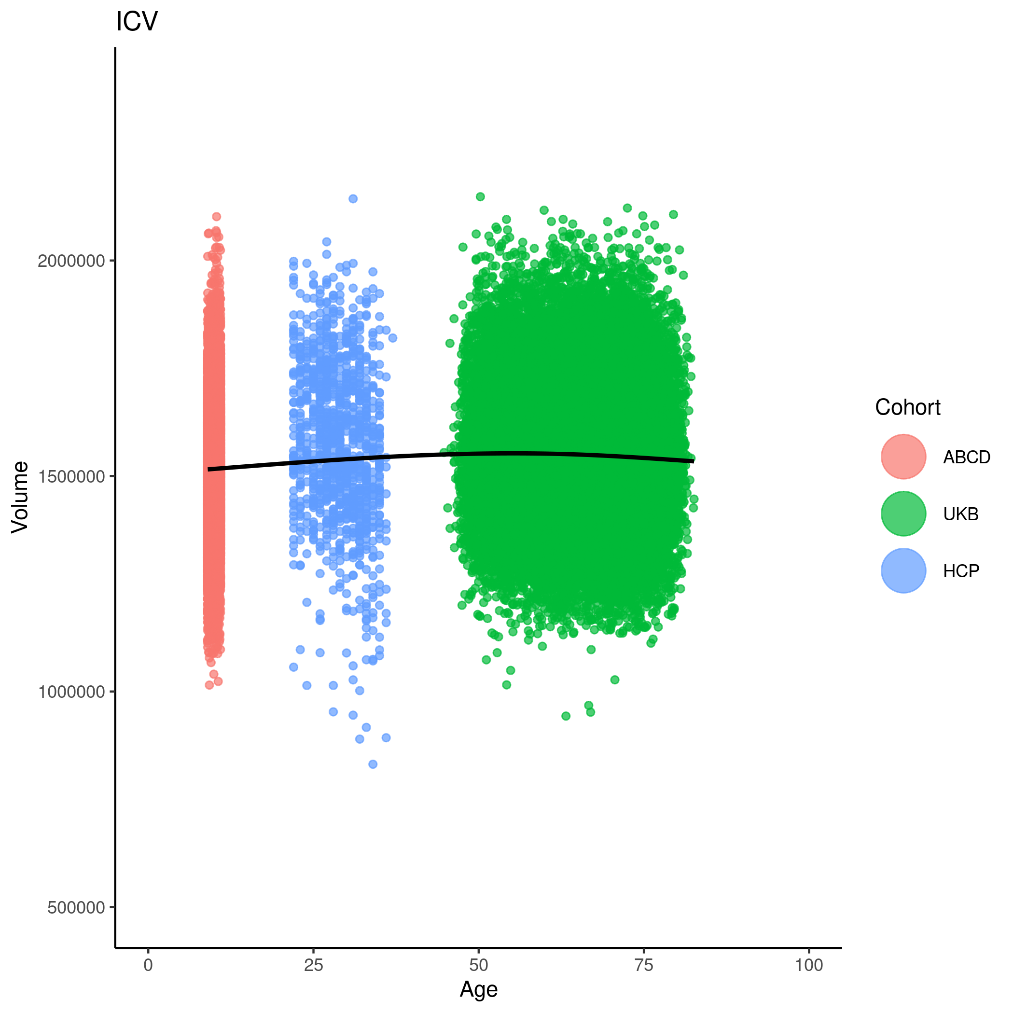 | 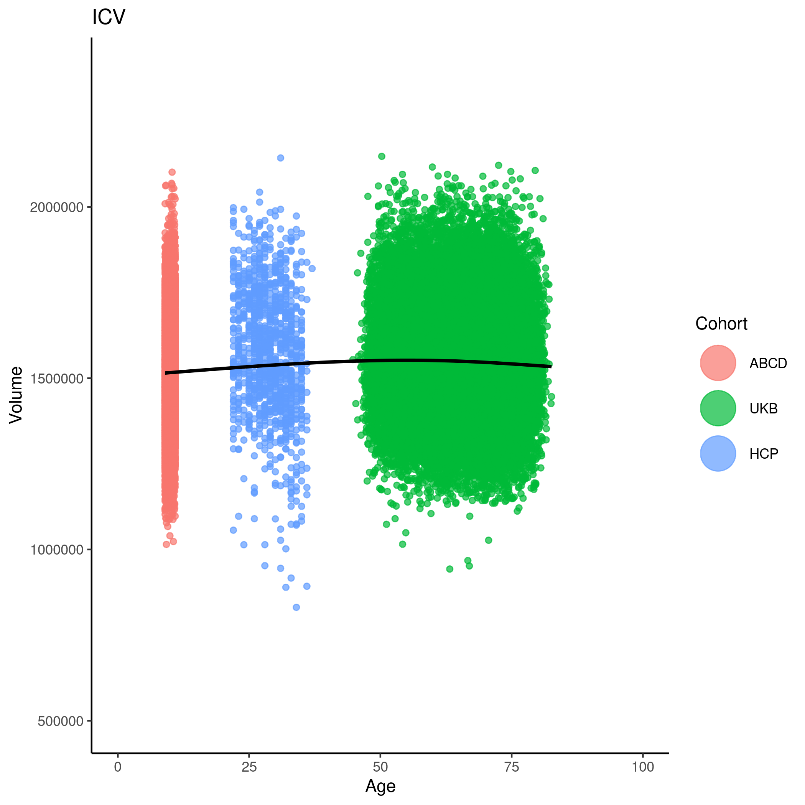 |

**Supplementary Figure 4. Age-related differences in GM and ICV.** Individual data points are shown across upper panel: All Lifebrain cohorts (BASE-II from Germany, Betula from Sweden, CALM, Cam-Can and Whitehall from the UK, HUBU from Denmark, LCBC from Norway, NESDA from the Netherlands, and UB from Spain), and lower panel: ABCD and HCP from the U.S., and UKB from the UK,. The black line represents a smoothed (3 knots) average difference. All data after outlier removal are shown.

**Supplementary Table 1. Standard deviations for years of education across samples**

|  | **Sample** | **Standard deviation for years of education** |
| --- | --- | --- |
| **European studies** | BASE-II | 2.839083 |
|  | Betula | 4.133765 |
|  | CALM | -- |
|  | Cam-CAN | 4.005441 |
|  | HUBU | 2.084957 |
|  | LCBC-Adult | 2.769779 |
|  | LCBC-Dev | 3.285162 |
|  | NESDA | 3.197675 |
|  | UB | 4.097159 |
|  | UKB | 1.507716 |
|  | Whitehall | 3.078847 |
| **US studies** | ABCD | 1.194957 |
|  | HCP | 1.796044 |

| **Income and education** | |
| --- | --- |
| **Development and Adulthood** | **Europe and US** |
| 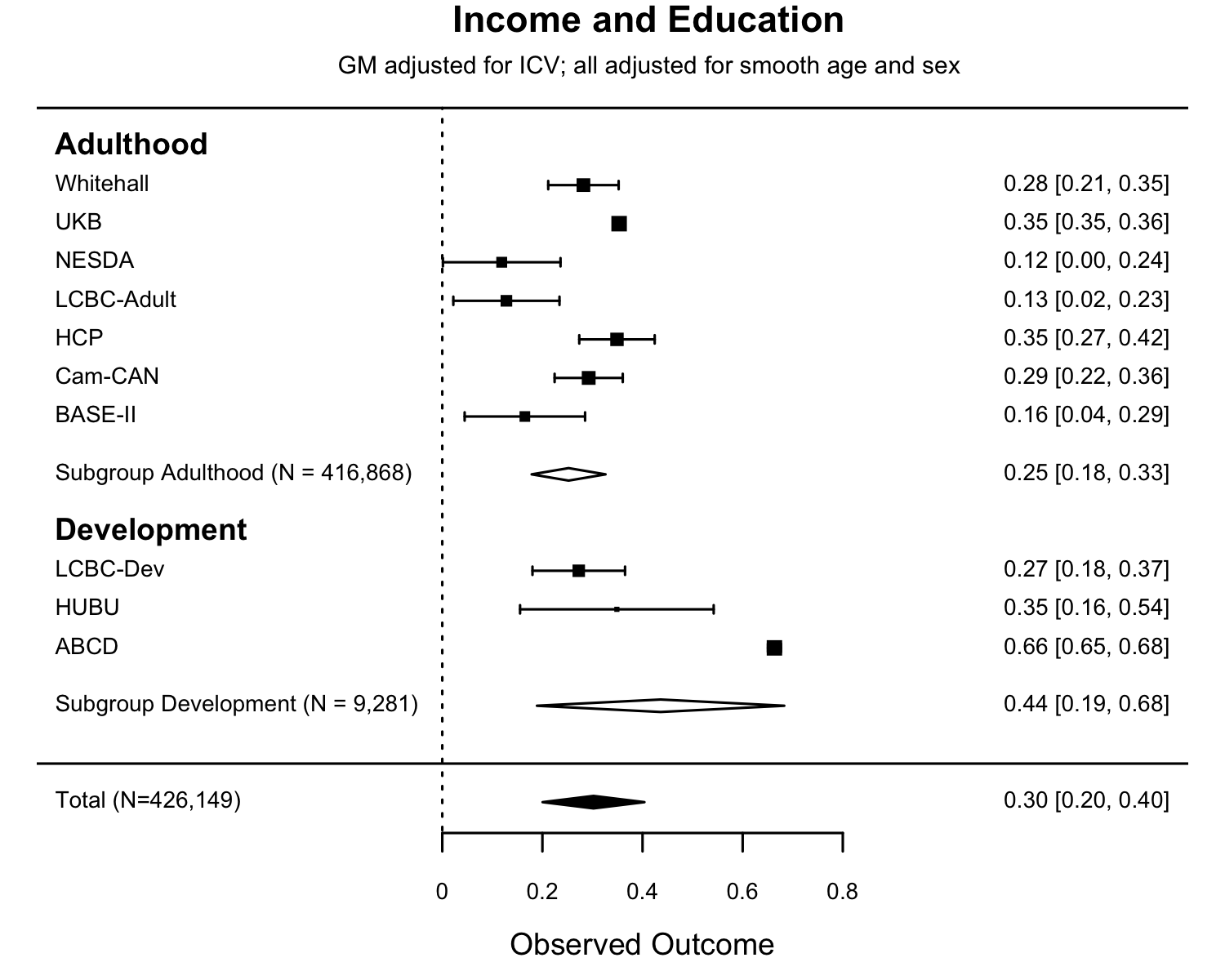 | 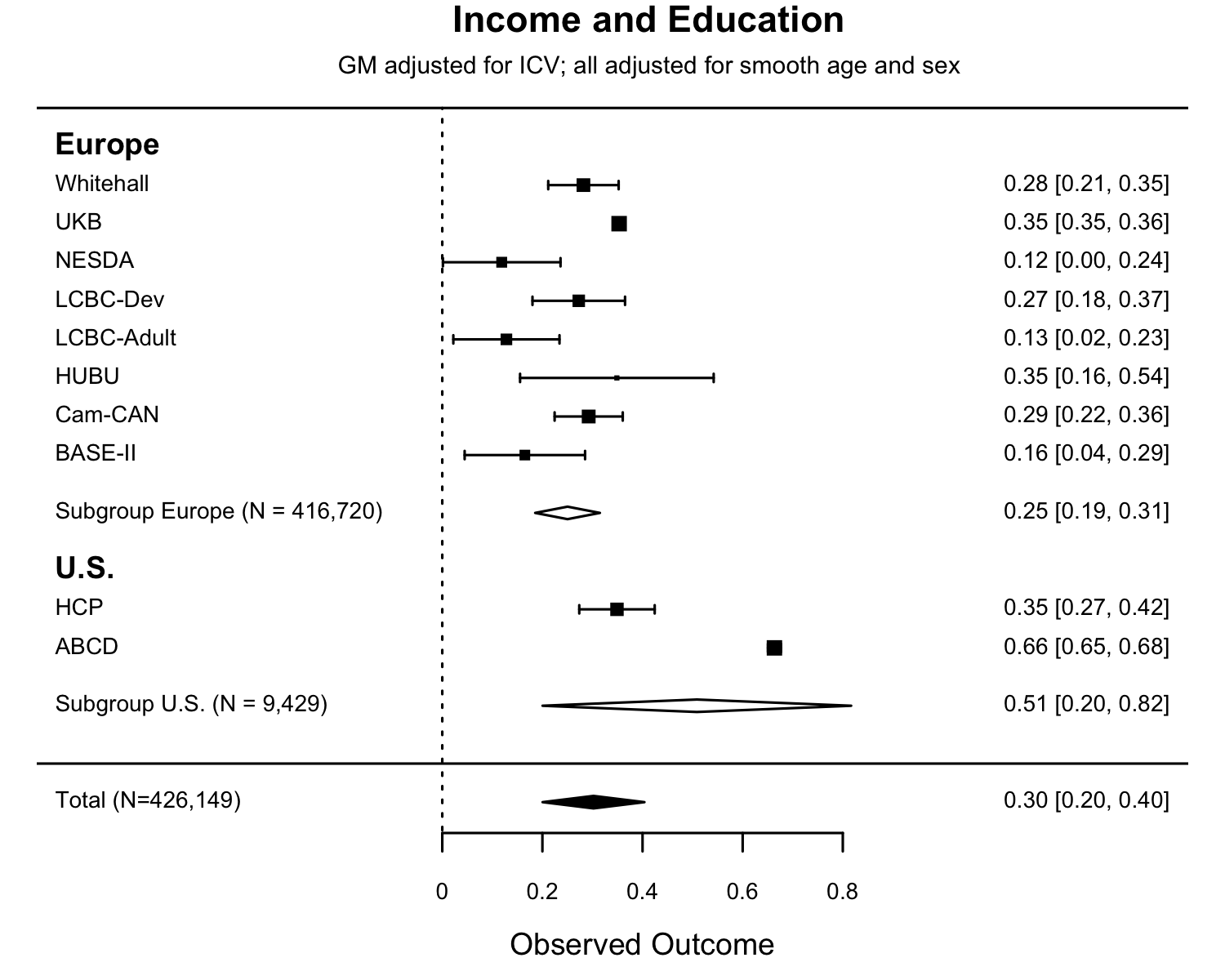 |

**Supplementary Figure 5** The associations of education and income for the cohorts. Forest plots show the individual observed effect sizes with corresponding 95% confidence intervals (CI). Diamonds represent the average correlation estimate and its 95% confidence interval with white diamonds representing the subgroup estimates and black diamonds the overall estimate Numeric values of the cohort-specific and meta-analytic estimates are given in the right column. The difference of association of income and education did not reach statistical significance for either Developmental vs. Adult cohorts (Z = -1.398, p= .1600), or European vs US cohorts (Z= -1.608, p= 0.1100).

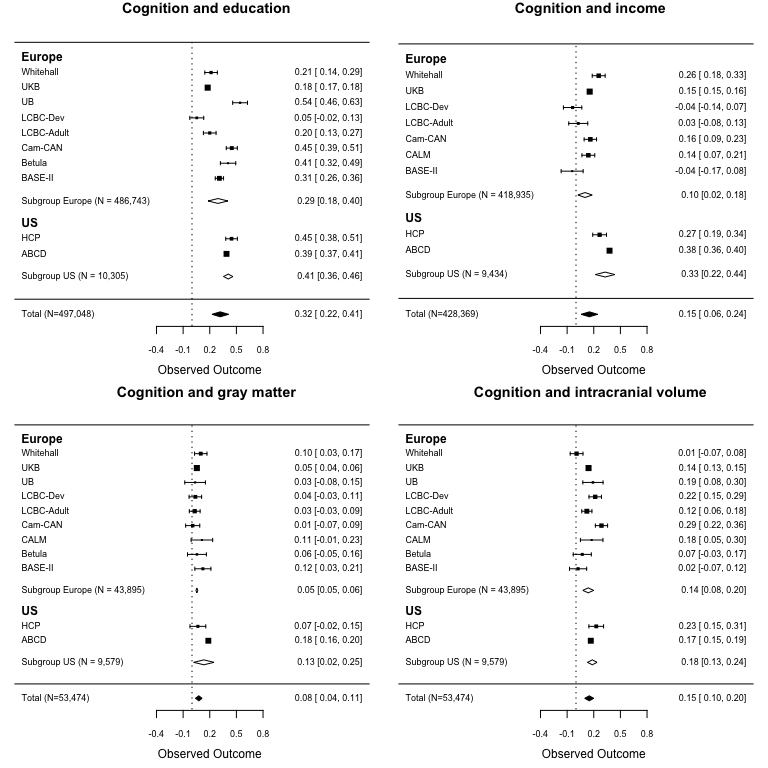

**Supplementary Figure 6.** The associations of cognition (general cognitive ability scores) with education, income, grey matter (GM) and intracranial volume (ICV), grouped by European vs US origin of cohorts. Forest plots show the individual observed effect sizes with corresponding 95% confidence intervals (CI). Diamonds represents the average correlation estimate and its 95% confidence interval with white diamonds representing the subgroup estimates and black diamonds the overall estimate. Numeric values of the cohort-specific and meta-analytic estimates are given in the right column. There were no significant differences in the associations of cognition with either education (Z= -1.831, p= 0.067), GM (Z =-.1.341, p = .1800), or ICV (Z =-.1.104, p = .2700) between European and US cohorts, but the association of cognition and income was stronger in the US (Z = -3.276, p= 0.0011).

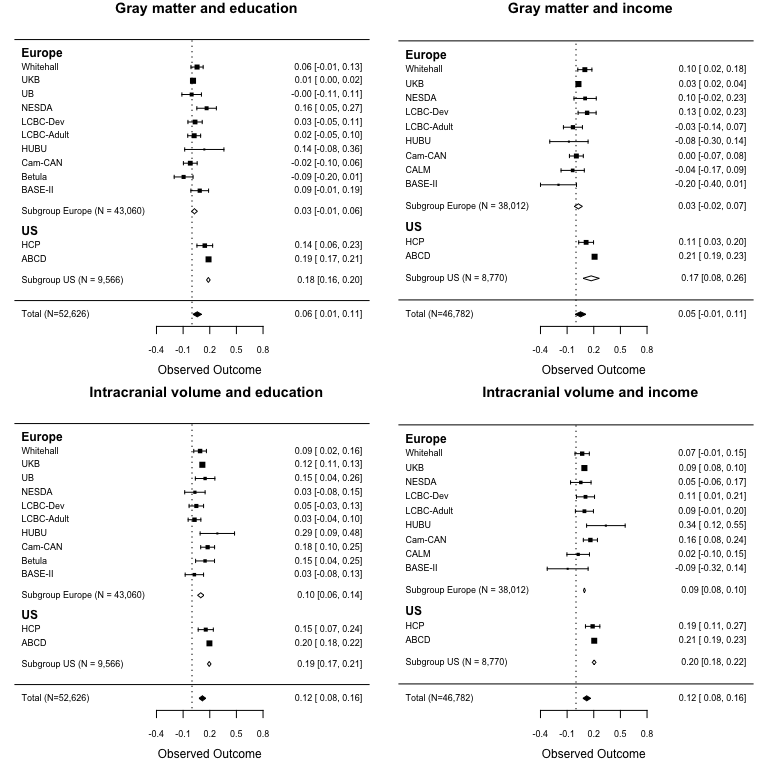

**Supplementary Figure 7.** The associations of grey matter (GM) and intracranial volume (ICV) to education and ICV, grouped by European vs US origin of cohorts. Forest plots show the individual observed effect sizes with corresponding 95% confidence intervals (CI). Diamonds represents the average correlation estimate and its 95% confidence interval with white diamonds representing the subgroup estimates and black diamonds the overall estimate Numeric values of the cohort-specific and meta-analytic estimates are given in the right column. There was a significantly greater association of GM adjusted for ICV with education (Z= -7.978, p< 0.0001) as well as income (Z= -2.763, p= 0.0057) in US than European cohorts. The association between ICV and income was also significantly greater (Z= -9.583, p < .0001) in US than European cohorts, and the same held for the association of ICV and education (Z= -4.528, p< .0001).

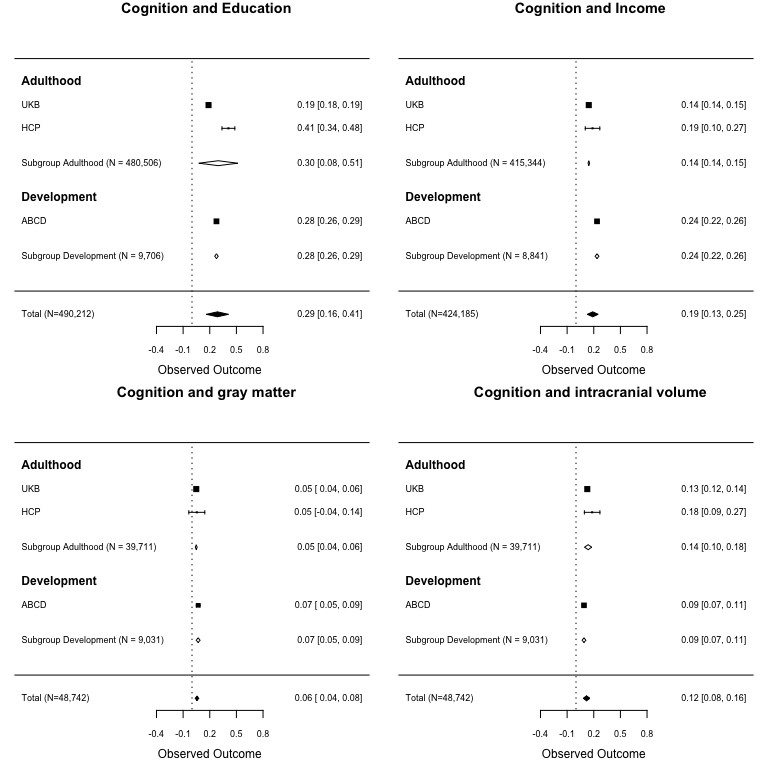

**Supplementary Figure 8** The associations of cognition (general cognitive ability scores) with education, income, grey matter (GM) and intracranial volume (ICV) when adjusted for reported ethnicity in addition to age and sex (compare to Figure 2). Forest plots show the individual observed effect sizes with corresponding 95% confidence intervals (CI). Diamonds represent the average correlation estimate and its 95% confidence interval with white diamonds representing the subgroup estimates and black diamonds the overall estimate. Numeric values of the cohort-specific and meta-analytic estimates are given in the right column.

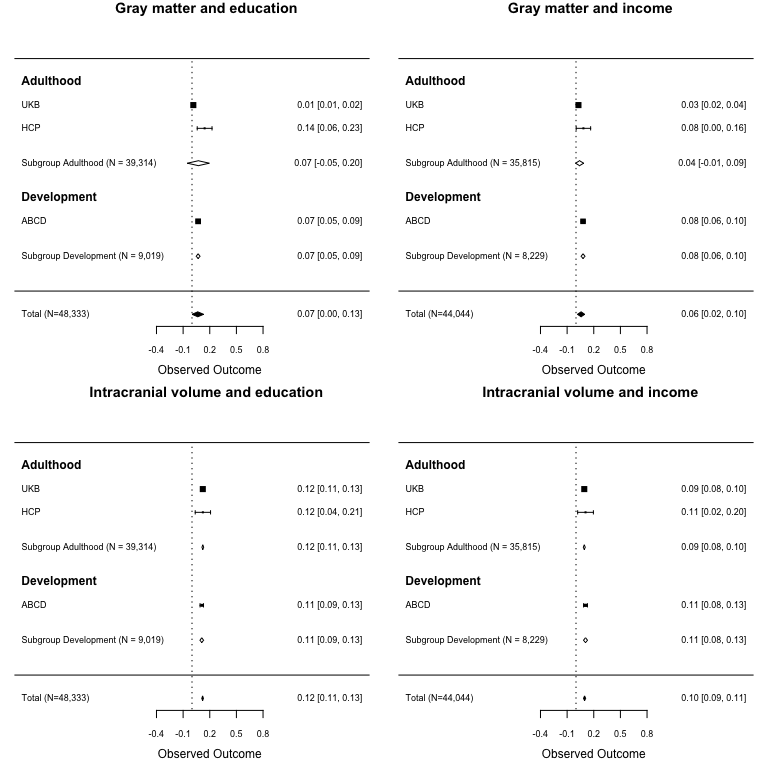

**Supplementary Figure 9** The associations of grey matter (GM) and intracranial volume (ICV) to education and ICV when adjusted for reported ethnicity in addition to age and sex (compare to Figure 3). Forest plots show the individual observed effect sizes with corresponding 95% confidence intervals (CI). Diamonds represents the average correlation estimate and its 95% confidence interval with white diamonds representing the subgroup estimates and black diamonds the overall estimate Numeric values of the cohort-specific and meta-analytic estimates are given in the right column.

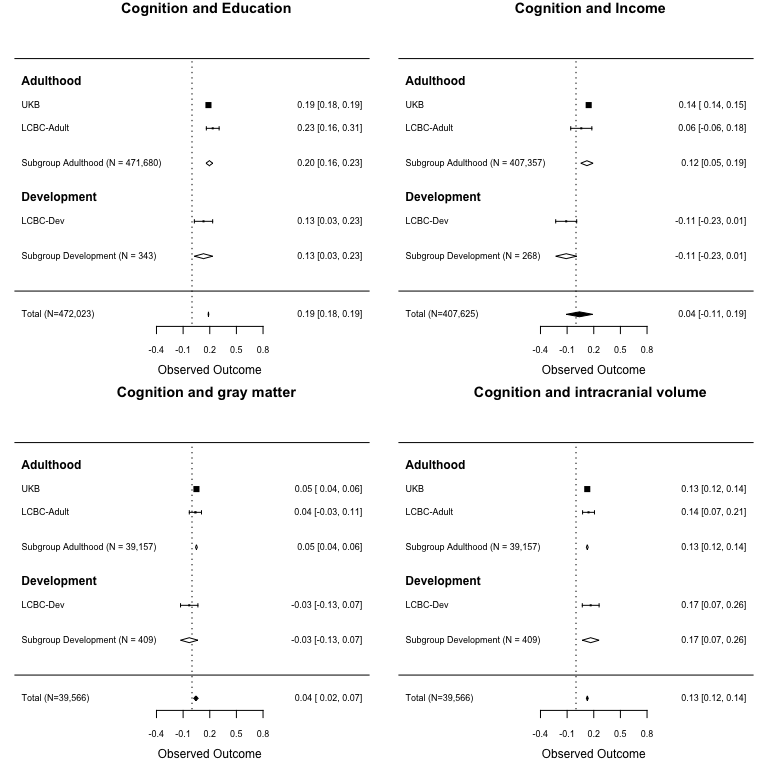

**Supplementary Figure 10** The associations of cognition (general cognitive ability scores) with education, income, grey matter (GM) and intracranial volume (ICV) when adjusted for genetic ancestry factors in addition to age and sex (compare to Figure 2). Forest plots show the individual observed effect sizes with corresponding 95% confidence intervals (CI). Diamonds represent the average correlation estimate and its 95% confidence interval with white diamonds representing the subgroup estimates and black diamonds the overall estimate. Numeric values of the cohort-specific and meta-analytic estimates are given in the right column.

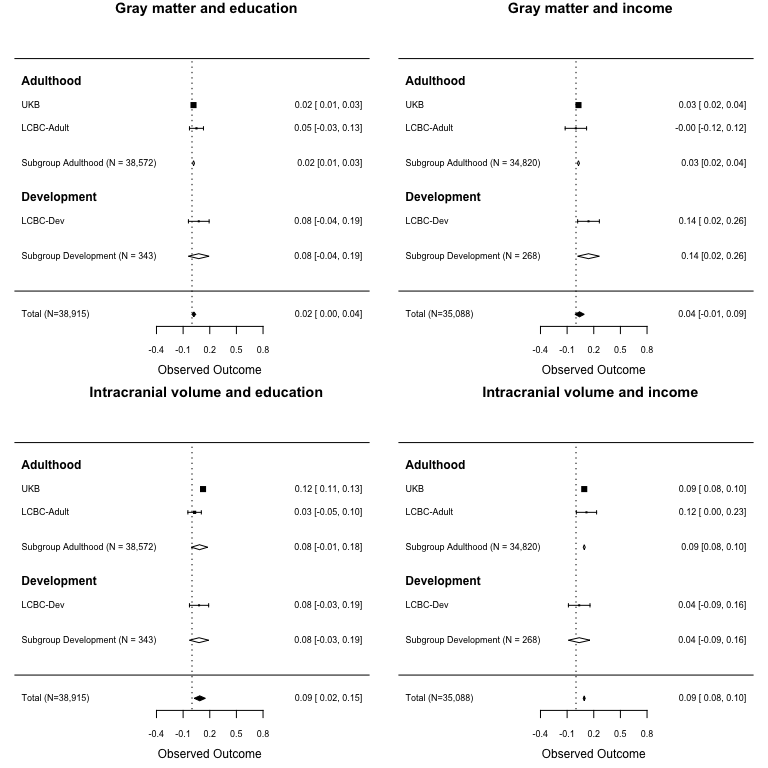
**Supplementary Figure 11** The associations of grey matter (GM) and intracranial volume (ICV) to education and ICV when adjusted for genetic ancestry factors in addition to age and sex (compare to Figure 3). Forest plots show the individual observed effect sizes with corresponding 95% confidence intervals (CI). Diamonds represents the average correlation estimate and its 95% confidence interval with white diamonds representing the subgroup estimates and black diamonds the overall estimate Numeric values of the cohort-specific and meta-analytic estimates are given in the right column.

| **Education and GM adjusted for ICV**  **vs Education and ICV** | **Income and GM adjusted for ICV**  **vs Income and ICV** |
| --- | --- |
| 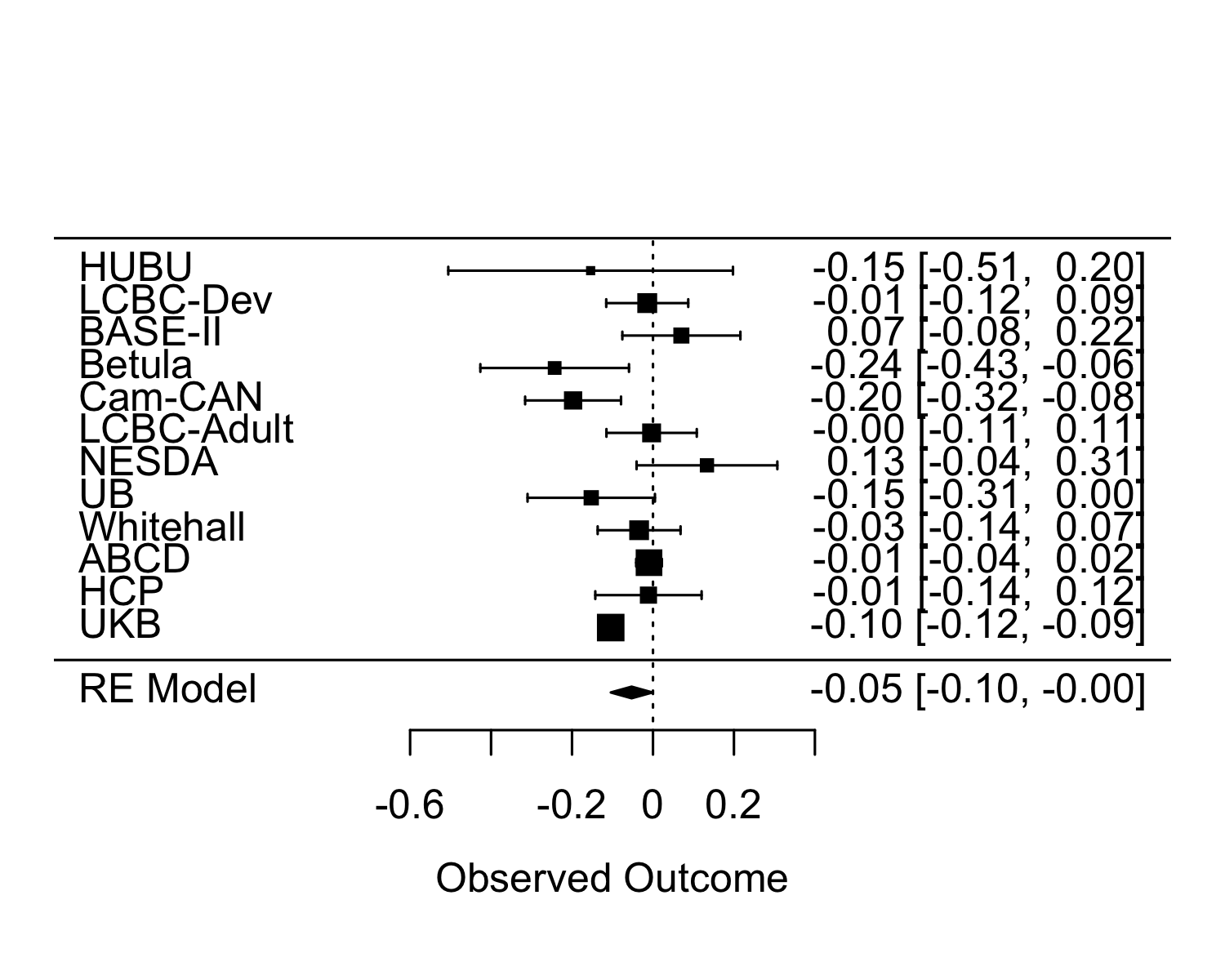 | 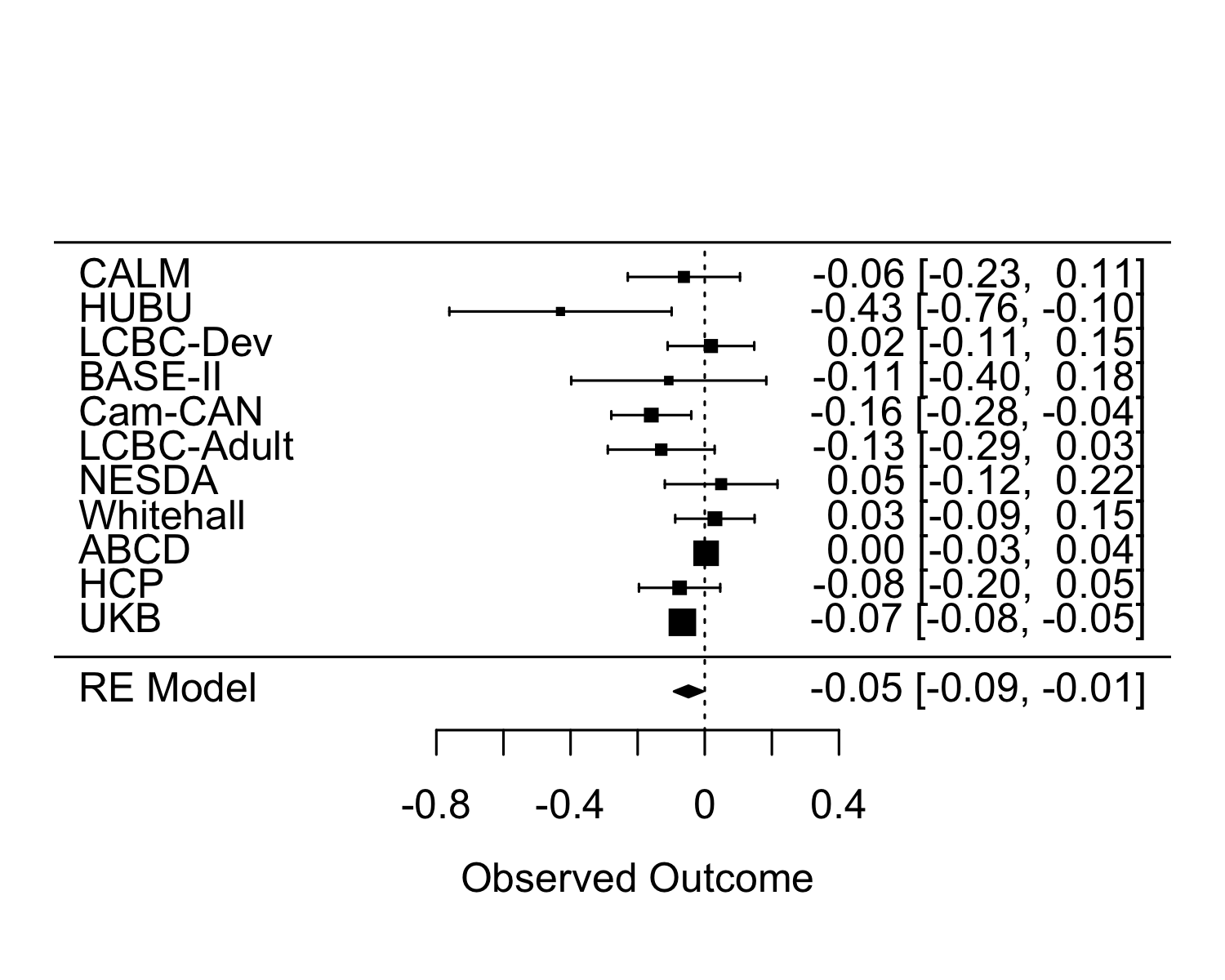 |
| Meta-analytic point estimate: -0.05275574  p= 0.04405358 | Meta-analytic point estimate: -0.04856496  p= 0.02700579 |
| **Cognition and GM adjusted for ICV**  **vs Cognition and ICV** | **Education and Cognition**  **vs. Income and Cognition** |
| 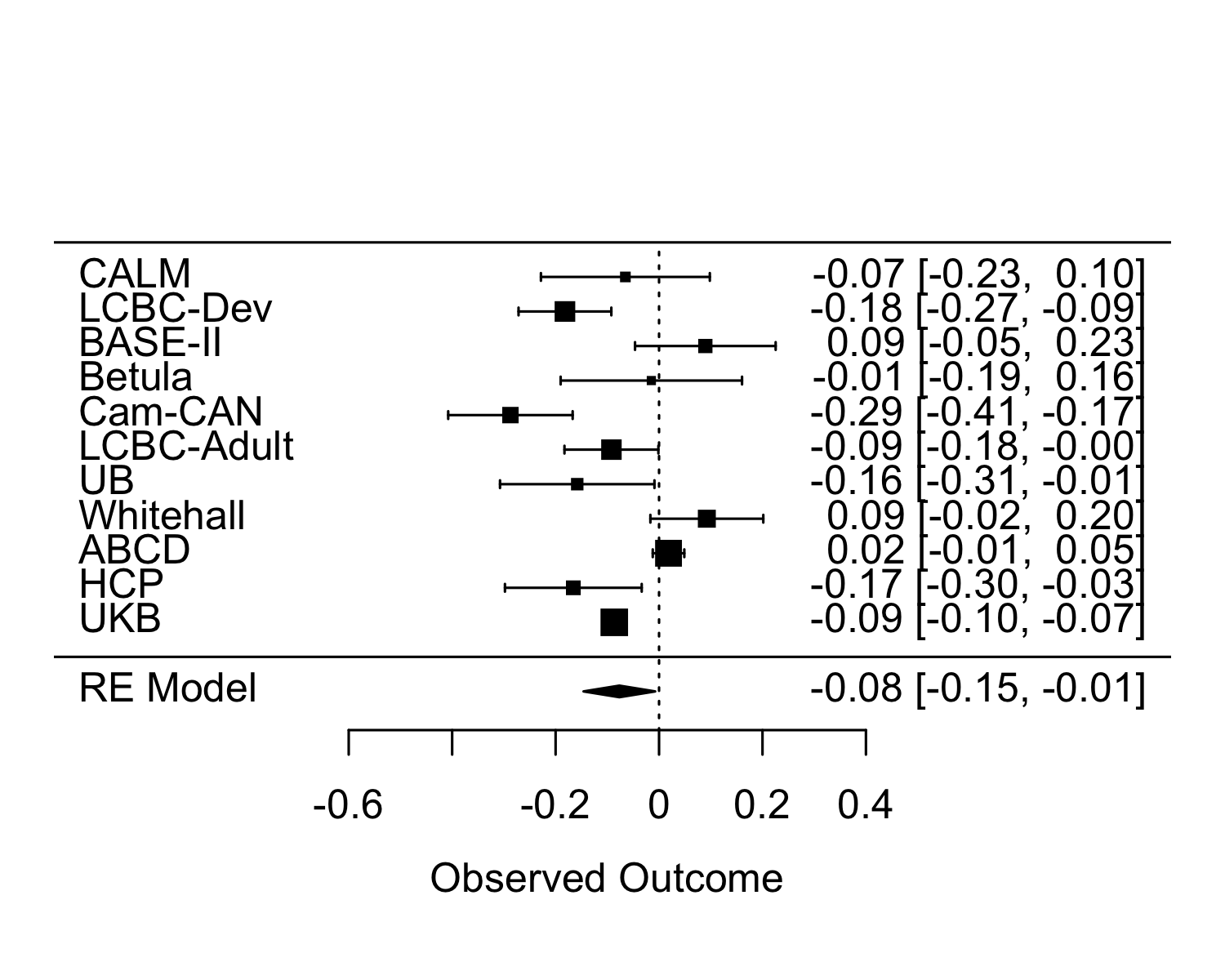 | 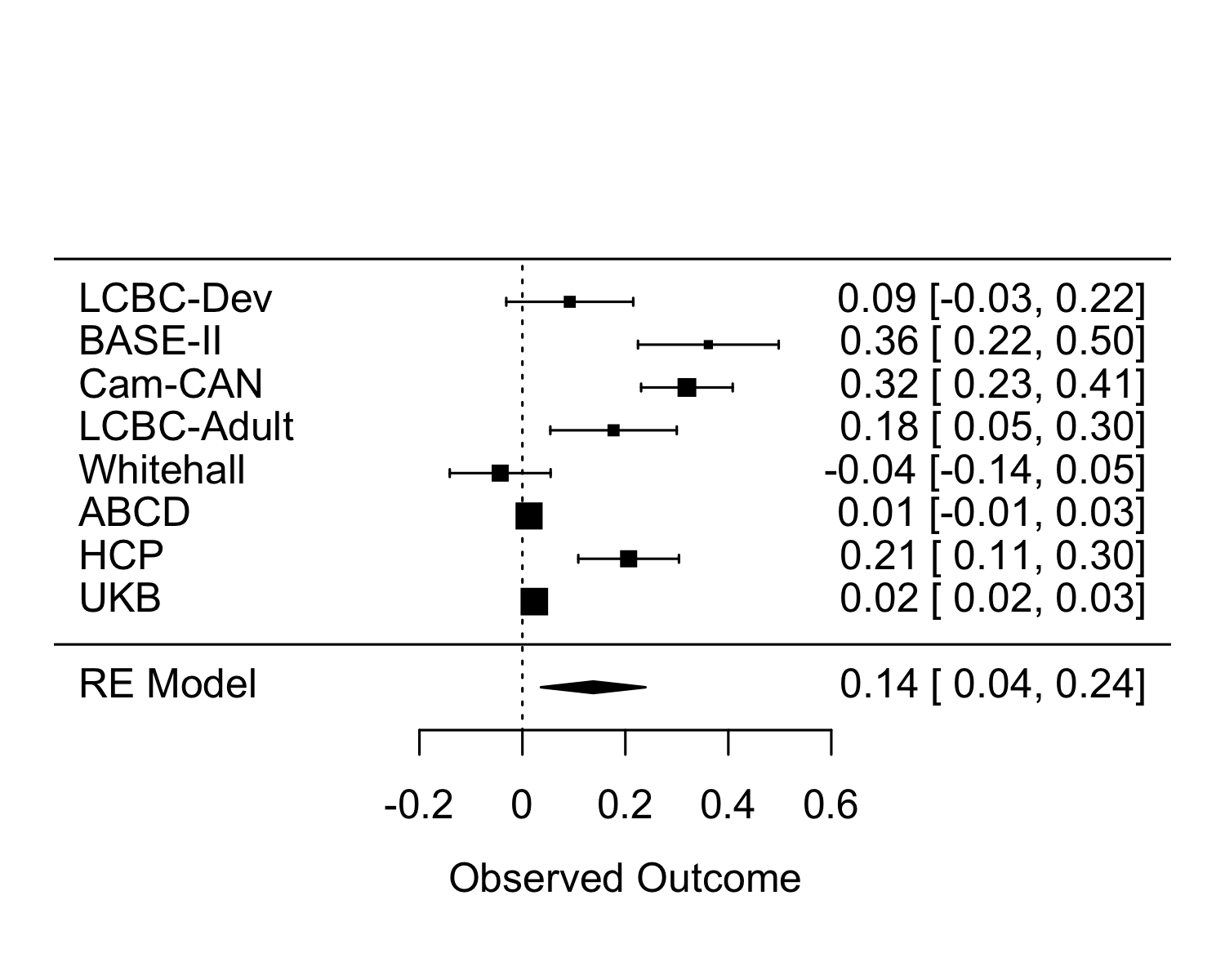 |
| Meta-analytic point estimate: -0.07674068  p= 0.02964948 | Meta-analytic point estimate: 0.1375985  p= 0.007894446 |

**Supplementary Figure** **12 Difference of relationships between SES and GM vs SES and ICV, Cognition and GM vs Cognition and ICV, Cognition and Education vs Cognition and Income**

Overall education was more related to ICV than to GM. Cognition was also more related to ICV than GM, and cognition was more related to education than to income. Left upper panel: We tested the difference of 1) correlation of education and GM (adjusted for ICV) with both being adjusted for smooth age and sex, 2) correlation of education and ICV with both being adjusted for smooth age and sex. Positive values mean that the correlation with GM is larger than the one with ICV. Right upper panel: We tested the difference of 1) correlation of income and GM (adjusted for ICV) with both being adjusted for smooth age and sex, 2) correlation of income and ICV with both being adjusted for smooth age and sex. Positive values mean that the correlation with GM is larger than the one with ICV. Left lower panel: We tested the difference of: 1) correlation of cognition and GM (adjusted for ICV) with both being adjusted for smooth age and sex, 2) correlation of cognition and ICV with both being adjusted for smooth age and sex. Positive values mean that the correlation with GM is larger than the one with ICV. Right lower panel: we tested the difference of 1) correlation of cognition and education with both being adjusted for smooth age and sex, 2) correlation of cognition and income with both being adjusted for smooth age and sex. Positive values mean that the correlation with education is larger than the one with income.

| **GCA-Education** | **GCA-Income** |
| --- | --- |
| **GM+ICV ‘mediates’ GCA <-> Education** | **GM+ICV ‘mediates’ Cognition <-> Income** |
| 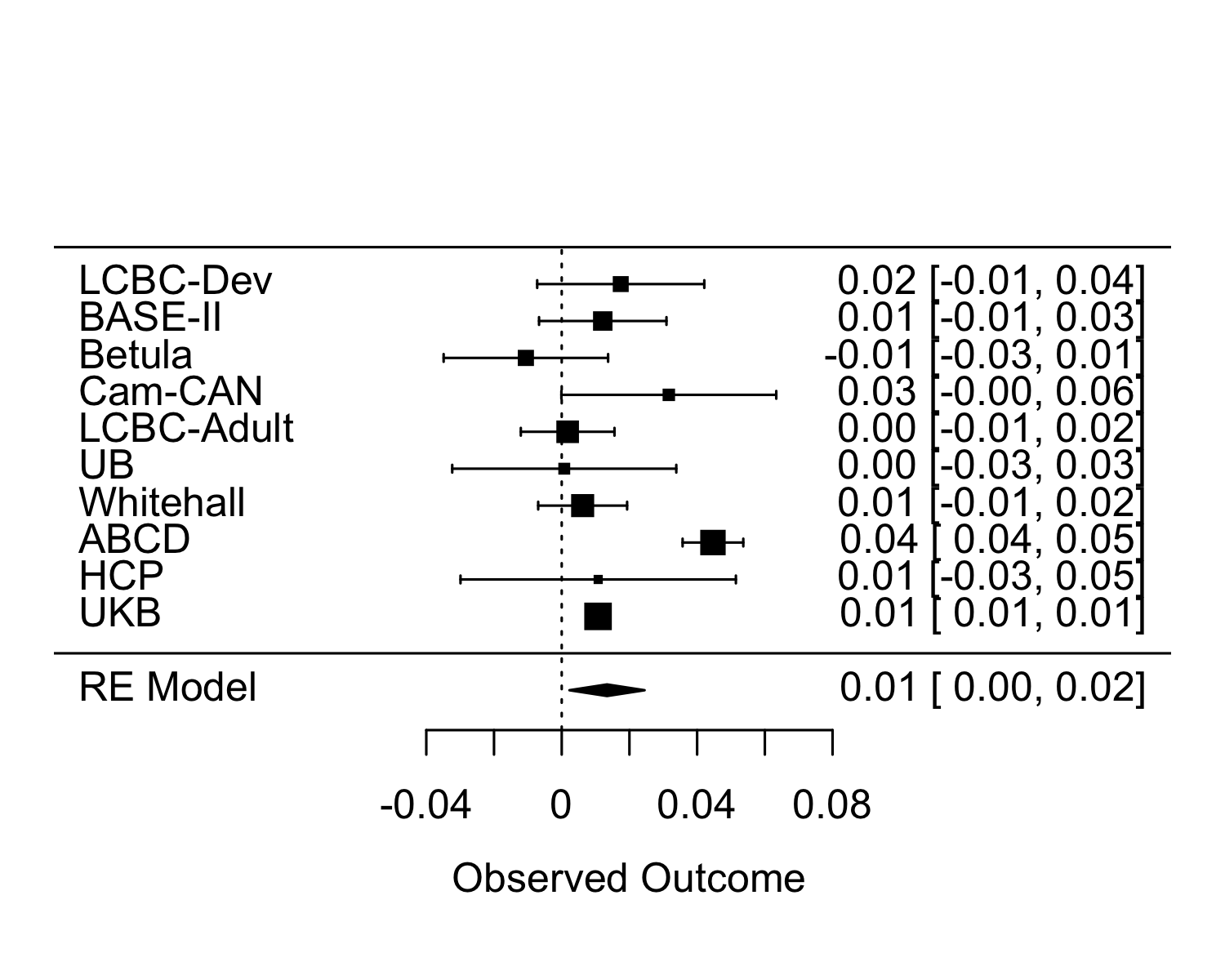 | 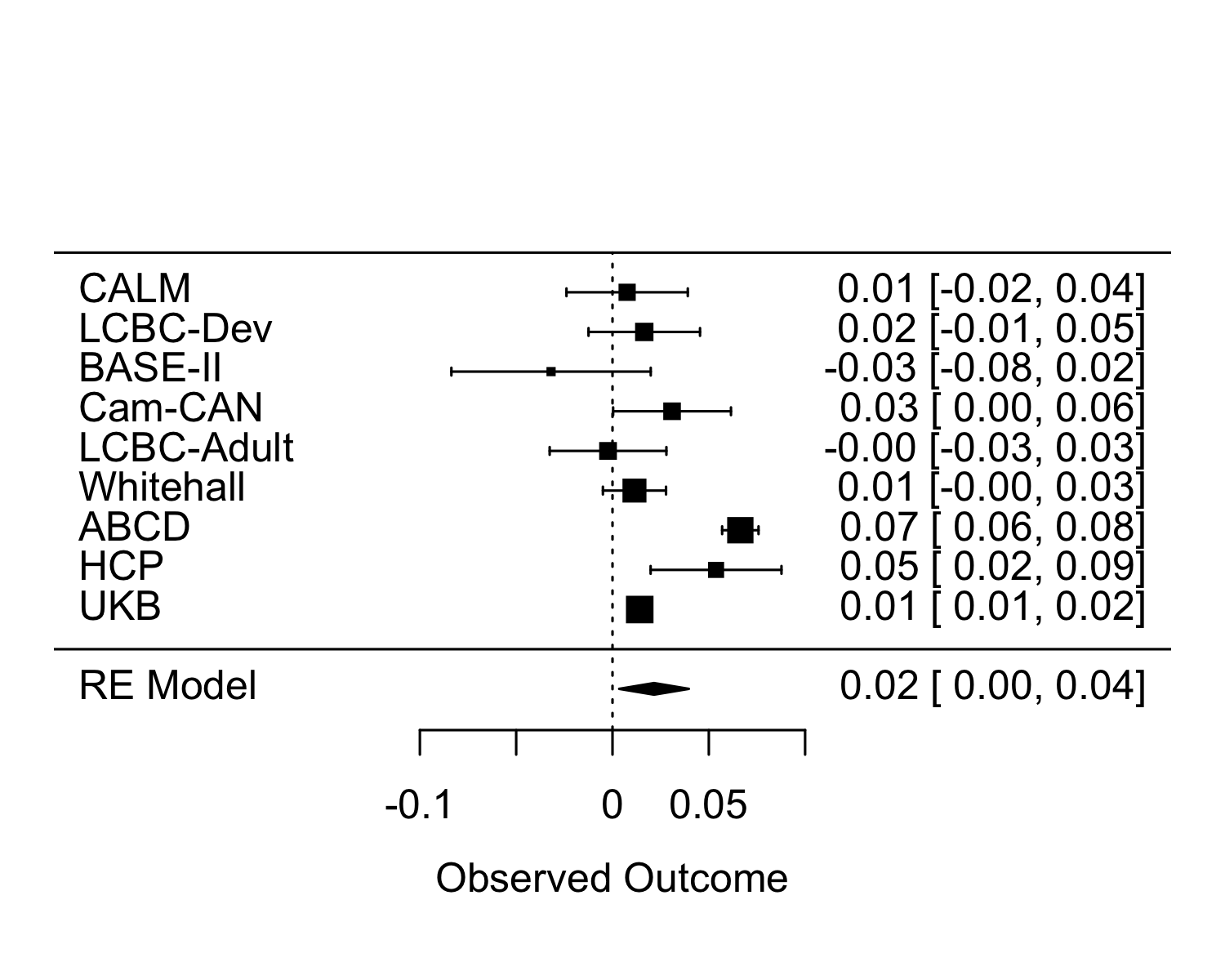 |
| Meta-analytic point estimate: 0.01338134 p= 0.01806167 | Meta-analytic point estimate: 0.02155208  p= 0.01940606 |
| **ICV ‘mediates’ GCA <-> Education** | **ICV ‘mediates’ GCA <-> Income** |
| 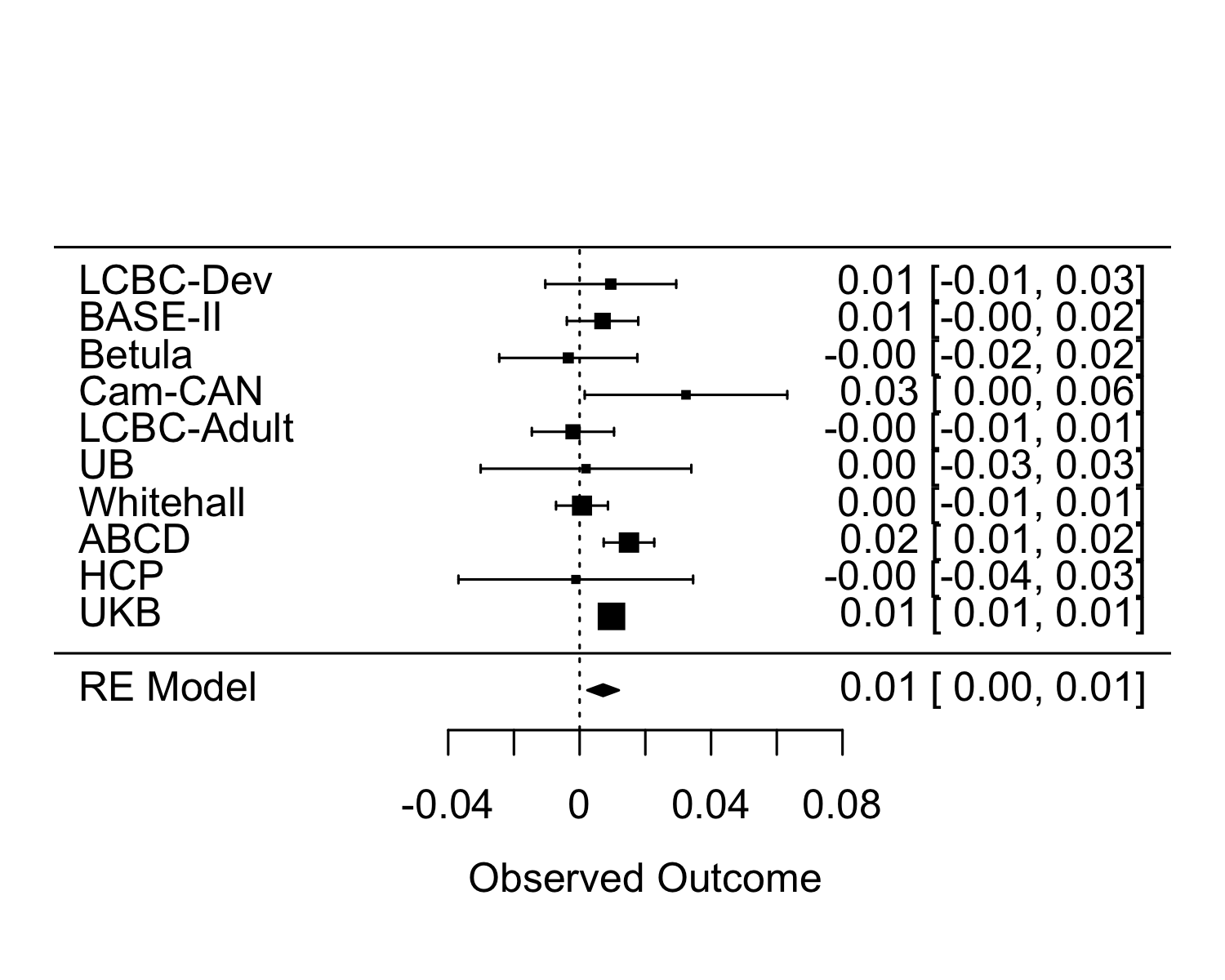 | 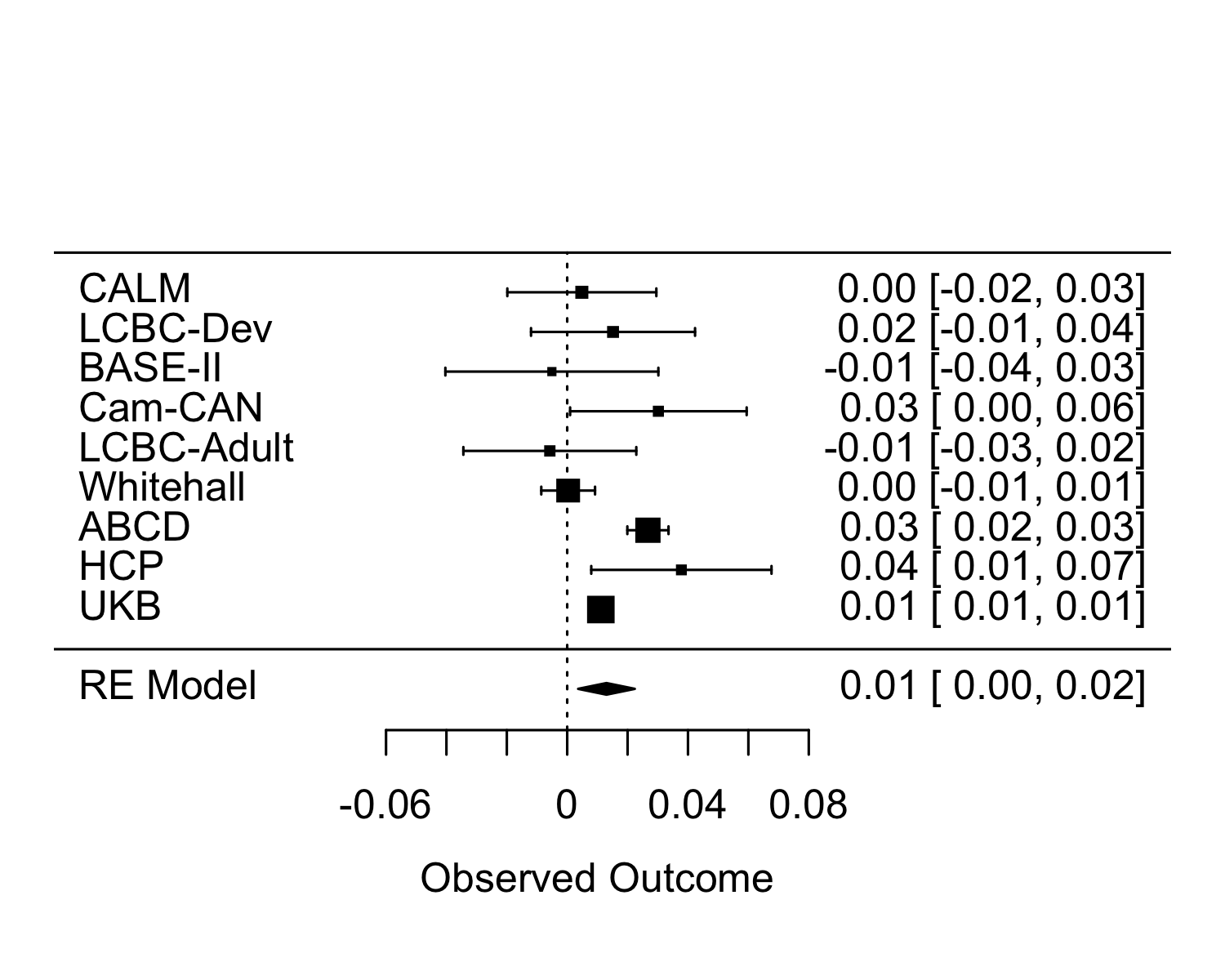 |
| Meta-analytic point estimate: 0.007142736  p= 0.00342541 | Meta-analytic point estimate: 0.01300417  p= 0.006502333 |
| **GM ‘mediates’ GCA <-> Education** | **GM ‘mediates’ GCA <-> Income** |
| 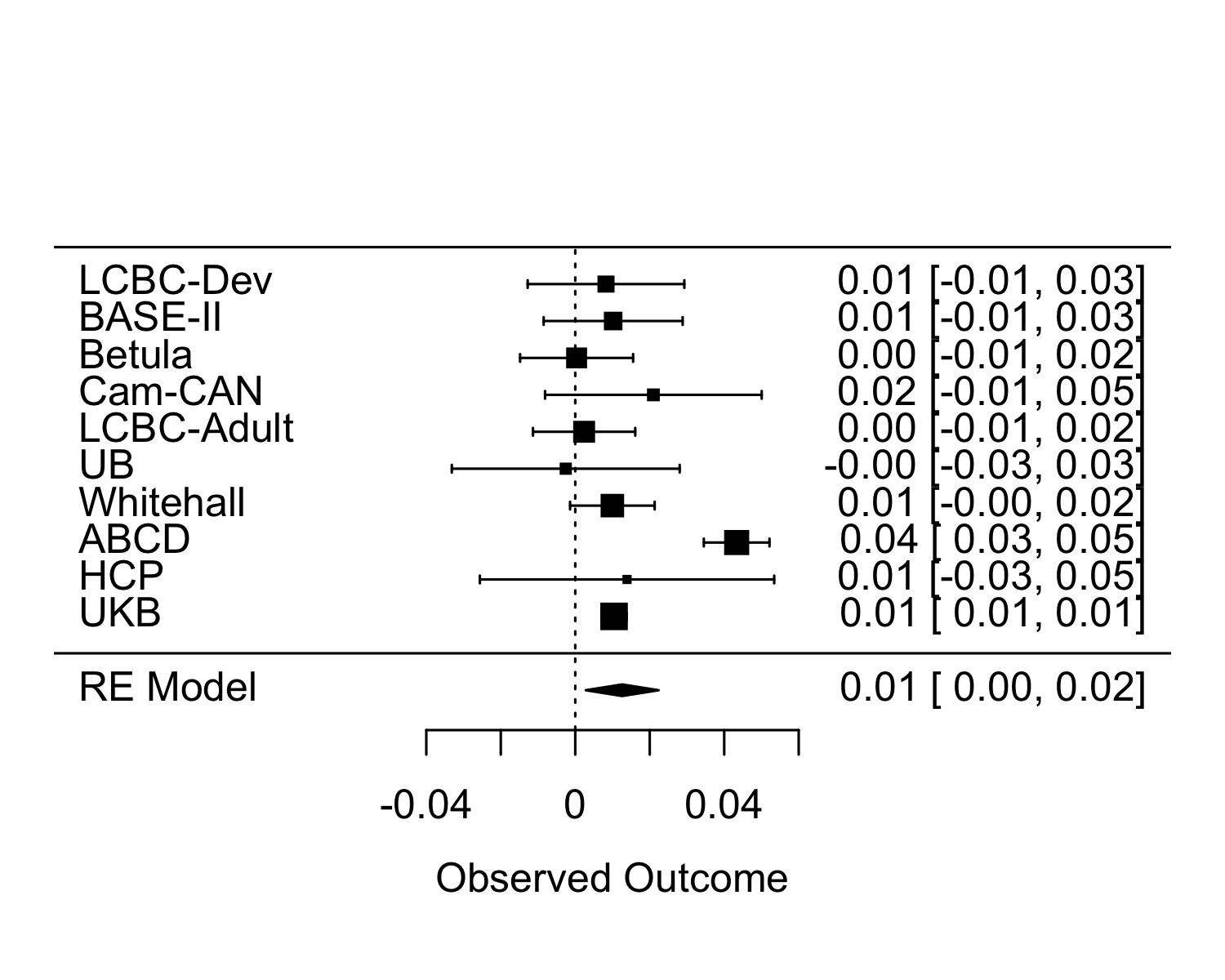 | 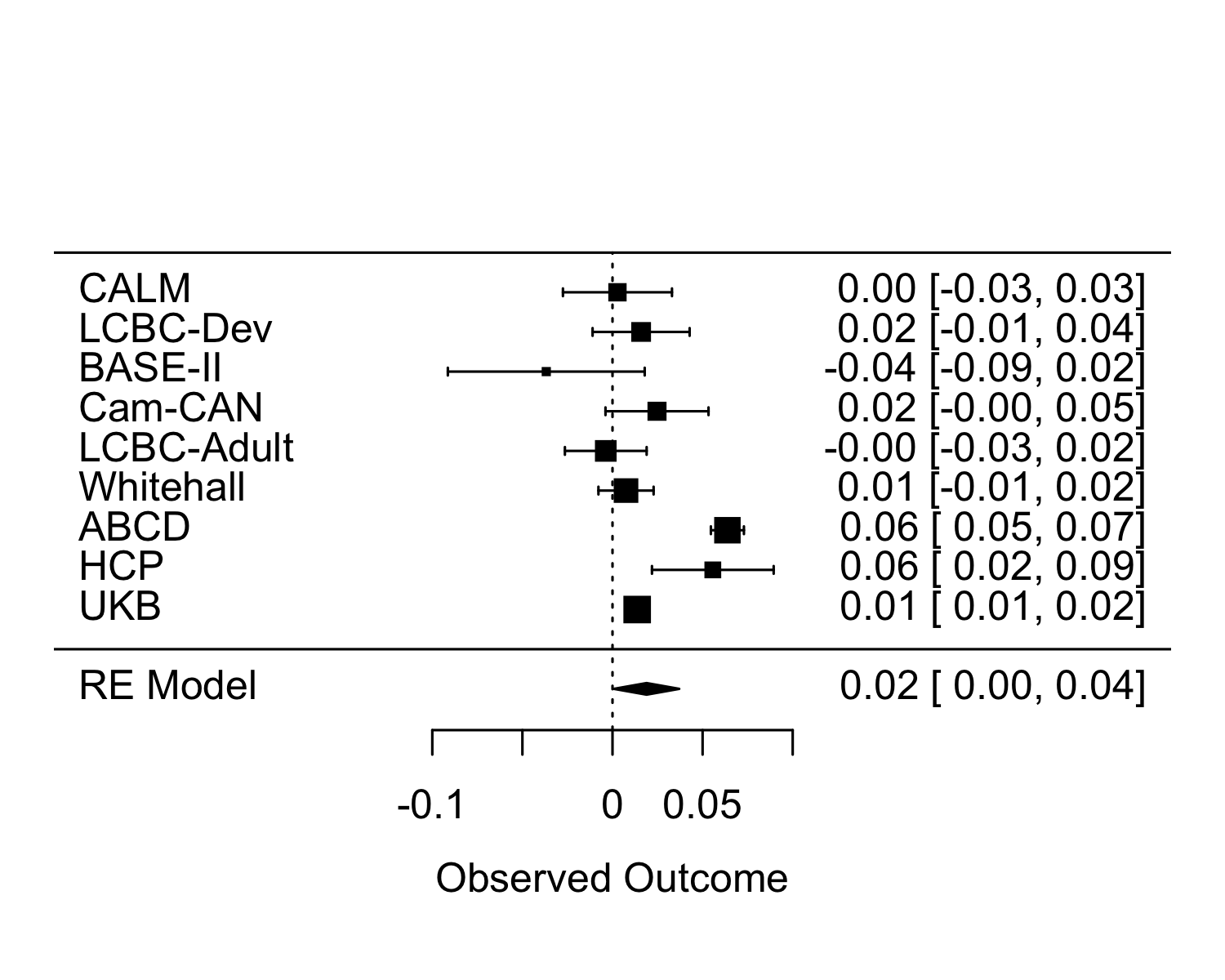 |
| Meta-analytic point estimate: 0.01259736  p= 0.01151603 | Meta-analytic point estimate: 0.01891117  p= 0.04055195 |

**Supplementary Figure 13 Difference of relationships between General Cognitive Ability (GCA) and SES when controlled for brain variables.**

We tested the difference of correlations of GCA and SES variables education (left panel) and income (right panel) with 1) both being adjusted for smooth age and sex 2) both being adjusted for smooth age and sex and either Intracranial Volume (ICV) and gray matter (GM) combined (upper panels), ICV alone (middle panels) or GM alone (lower panels). Positive values mean that the correlation unadjusted for brain values is larger than the adjusted one. As seen, overall, the correlations between GCA and SES were larger when not being adjusted for brain variables, and this difference was significant overall for relationships between education and GCA as well as income and GCA, indicating that variance in the brain variables explained part of the relationships. There is considerable variance across cohorts, in the extent to which these variables altered the correlations.

**Supplementary methods**

**Supplementary sample information**

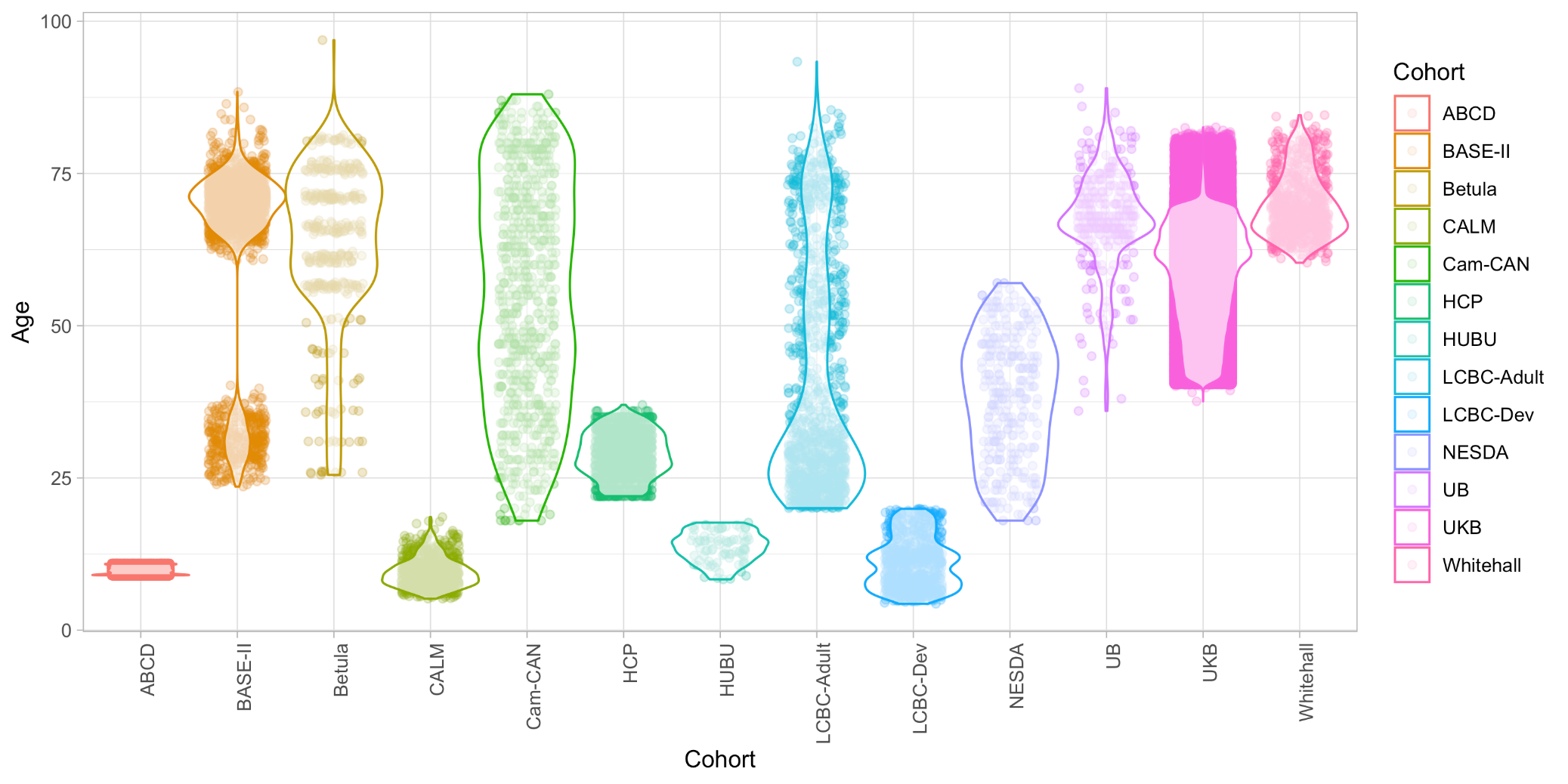

**Supplementary Figure 14** Violin and dot plots of age ranges covered and age distributions in each cohort.

**Sample recruitment, inclusion and exclusion criteria, demographics and measures**

**BASE II**

*Population, recruitment, inclusion/exclusion criteria and general description of study*

Participants of the Berlin Aging Study II (BASE II) were community-dwelling older adults recruited from the greater Berlin metropolitan area through advertisements in newspapers and public areas (for cohort characteristics and additional details, see ^5,6^). The baseline sample comprised 2200 participants. Of these, 1600 were older adults aged 61–88 years (mean age 71.5, SD 3.89; 793 female), and 600 were younger adults aged 24–40 years (mean age 31.1, SD 3.38; 247 female). Participants were invited to a medical exam consisting of a 2-day protocol, and two cognitive testing sessions scheduled 1 week apart, and were tested in small groups (e.g. about 6 participants per group) on a comprehensive cognitive battery that covers key cognitive abilities measured by 21 tasks. Each session lasted about 3.5 h. 1828 had valid data to be included in the present analyses.

MR Sample (TP1): After completion of the cognitive examination of BASE-II, eligible participants were invited to take part in one MRI session within a time window of 2–4 weeks after cognitive testing, consisting of 341 older adults aged 61–82 years (mean age 70.1, SD = 3.89; 131 female) and 103 younger adults (mean age 31.4, SD = 3.7; 39 female). In total, 414 of these had valid data to be included in the present MR analyses. MR scans and cognitive scores were obtained 2012-2013. A subsample of the MR sample was later re-invited for follow-up, but only baseline data are included in the present analysis. The different elements of the study were approved by the ethics committees of the Max Planck Institute for Human Development, the Charité University ethics committee and by the ethics committees of The German Association for Psychology (DGPs). Participants signed written informed consent and received monetary compensation for their participation in BASE-II and the MRI study. All experiments were performed in accordance with relevant guidelines and regulations.

*Inclusion/exclusion criteria/ screening*

Inclusion criteria for taking part in this study were age between 20 and 35 or 60 and 80 years, apparently healthy. Exclusion criteria were untreated diabetes and hypertension; prior stroke, head injuries or brain surgery; psychiatric illness; major depression; dementia with a score < 24 on the Mini-Mental State Examination. To that end, none of the participants took medication that might affect memory function or had a history of head injuries, medical (e.g., heart attack), neurological (e.g., epilepsy), or psychiatric disorders (e.g., depression). All participants reported normal or corrected to normal vision, were right-handed, and scored over 27 on the Mini-Mental Status Examination.

*Variables used for education, income and general cognitive ability (GCA)*

Education was measured as number of years spent in formal schooling, also correcting for east/west German educational systems. Income was measured by questionnaire items asking for last month´s income (including any salary/pension after tax, except special payments such as e.g. vacation allowance). The current measure of Fluid Cognitive Ability on which the GCA measure used in analyses is based, are Practical Problems, Figural Analogies, Letter Series, as described in ^7^.

**Betula**

*Population, recruitment and general description of study/ procedures*

In the Betula longitudinal study on aging, memory and dementia, population-based sampling of healthy middle-aged and older adults was used for recruitment. Detailed recruitment procedures are found in ^8,9^. For the current analyses, the MRI subsample of the study is used. Participation in the neuroimaging study was offered to all participants who had remained in the study and completed cognitive testing at the 5th Betula test wave in 2008-2009, and 376 participants underwent structural and functional MRI in 2009-2010.

*Inclusion/exclusion criteria/ screening*

Exclusion criteria were severe visual or auditory handicaps, intellectual or developmental disabilities, suspected dementia, having a mother tongue other than Swedish, MRI contraindications, neurological disorders, or visual/motor deficits that could interfere with fMRI data collection, MMSE <24, brain or head surgery, and substantial brain anatomical deviations. Eight participants were excluded completely post scanning due to discovered neurological conditions (Schizophrenia, Multiple Sclerosis, Parkinson’s Disease, Hydrocephalus, Alcoholism, and dementia), and an additional two participants were excluded due to MMSE scores below 24. In addition, for 29 participants MRI data only was excluded due to anatomical deviations (subdural hematoma, localized loss of brain tissue, subcortical atrophy, and previous brain or head surgery (n=2)), movement artifacts (n=21), or FreeSurfer processing failures (n=3). Three individuals had missing T1 images due to incomplete acquisition. Thus, the final data set comprised 366 participants, of which 364 provided cognitive and SES data, 332 SES and MRI volumetric data, while 2 participants only provided MRI, age, and sex.

*Variables used for education, income and general cognitive ability (GCA)*

Age at MRI-scanning, reported with a one decimal precision, was used in the analyses. Self-reported years of education with a half-year precision was used as the measure of SES. Cognitive testing was performed 1-18 months prior to scanning (mean interval: 9 months). For cognitive variables, fluid ability was estimated from the Block Design test ^10^, and a measure of immediate free recall of 16 enacted verb-noun sentences. Crystallized abilities were estimated from a 30-item five-alternative forced choice vocabulary test, four measures of verbal fluency measured during one minute (generating words starting with the letter A, professions starting with the letter B, five-letter words starting with letter M, and words starting with F, without the letter A), as well as a 26-item general knowledge test. Exact testing procedures have been described in more detail elsewhere ^8^.

**CALM**

*Population, recruitment and general description of study/ procedures*

Participants were recruited based on referrals made for possible attention, memory, language, reading and/or mathematics problems ^11^. Participants with or without formal clinical diagnosis were referred to CALM. A subset of participants completed MRI scanning (n = 258 in the present study).

*Inclusion/exclusion criteria/ screening*

Exclusion criteria included known signiﬁcant and uncorrected problems in vision or hearing and a native language other than English.

*Variables used for education, income and general cognitive ability (GCA)*

For crystalized ability, Single Word reading and Spelling were used ^12^. For fluid reasoning, we used matrix reasoning ^13^ and Digit Recall (immediate serial recall of sequences of up to 9 spoken digits) ^12^. For income, the following question was asked:

“What is your total household income after tax? If you are financially independent, please tick your own income after tax. Please include any child support or benefits that you may receive.”, using the categories £0 - £9,999; £10,000 - £19,999; £20,000 - £29,999; £30,000 - £39,999; £40,000 - £49,999; £50,000 - £59,999; £60,000 - £69,999; £70,000 - £79,999; >£80,000; Prefer not to say.

**Cam-Can**

*Population, recruitment and general description of study/ procedures*

Recruitment was done by invitation letters based on the patient lists of general practitioners within the Cambridge City area. A population-based cohort of 3000 adults aged 18 or above was recruited to Stage 1 of the project, where they completed an interview including health and lifestyle questions, a core cognitive assessment, and a self-completed questionnaire of lifetime experiences and physical activity. Of those interviewed, ~700 participants aged 18-87 (100 per age decile) continued to Stage 2 where they undergo cognitive testing and provide measures of brain structure and function. A subset of ~250 adults returned for longitudinal follow-up data. The study is conducted in compliance with the Helsinki Declaration, and has been approved by the local ethics committee, Cambridgeshire 2 Research Ethics Committee (reference: 10/H0308/50).

*Inclusion/exclusion criteria/ screening*

General exclusion criteria: Term-time residents of colleges and universities, and participants whose Primary Care Physician feel are inappropriate to include. Exclusion criteria for the MRI part of the study: Not cognitively normal (MMSE < 24, memory defect, consent difficulties), communication difficulties (hearing problems [35db at 1000 Hz], insufficient English language, vision difficulties), medical problems by self-report of diagnosis (dementia diagnosis /Alzheimer’s Disease, Parkinson’s Disease, Motor Neurone disease, Multiple sclerosis, cancer, stroke, encephalitis, meningitis, epilepsy, head injury with serious results [coma, unconscious for >2 hours, skull fracture], recently diagnosed or uncontrolled high blood pressure, possible pregnancy, current psychiatric conditions [bipolar disorder, schizophrenia, psychosis]), mobility problems (restricted mobility which could prevent further participation, inability to walk 10 metres), substance abuse (past or current treatment for drug abuse, current drug usage), MRI/ MEG safety and comfort exclusions.

*Variables used for education, income and general cognitive ability (GCA)*

For fluid abilities, the standard form of the Cattell Culture Fair, Scale 2 Form A, was used ^14^. The Cattell test is a pen-and-paper test where the participant chooses a response on each trial from multiple choices, and records responses on an answer sheet. Our crystalized measure was the Spot The Word task ^15^. Spot the Word was time-limited to 5 minutes due to interview time restrictions but this was not indicated to the participants and interviewers could be flexible.

For income, the following question was asked: “What is the average total income before tax received by your household?” Participant chose one of the following categories: Less than £18,000; £18,000 to £30,999; £31,000 to £51,999; £52,000 to £100,000; Greater than £100,000; Prefer not to answer. For education, the following question was asked: “At what age did you complete your continuous full time education? Participants answered by entering a number, or “00” for “Never went to school”, “77” for “Do not know” or “88” for “No answer”.

**HUBU**

*Population, recruitment and general description of study/ procedures*

The present study included 86 typically-developing children and adolescents (49 girls, 37 boys) aged 8.2–17.4 years. Seventy-three participants were right-handed, 10 participants were left-handed and three participants were ambidextrous as assessed by the Edinburgh Handedness Inventory. 84 of these had valid MRI data. All children were enrolled in the longitudinal HUBU (“Hjernens Udvikling hos Børn og Unge” – in English: Brain maturation in children and adolescents) project designed to trace developmental changes conducted at the Danish Research Center for Magnetic Resonance. The total HUBU cohort include 95 typically-developing children (55 girls, 40 boys) and their families, who were recruited from three elementary schools in the Copenhagen suburban area in 2007, when the children were between seven to 13 years of age. Participants in the HUBU cohort have been assessed up to 12 times, with six months intervals for the first 10 assessments. In the present study, the first assessment with structural scans of good quality for each participant is included in the analyses. Prior to participation and after receiving oral and written explanation about the study aims and procedures, all children assented to partake in the study and informed written consent was obtained from the parents of all participants. The study was approved by the Ethical Committees of the Capital Region of Denmark (H-KF-01–131/03) and conducted in accordance with the Declaration of Helsinki. For more information on the HUBU cohort, see ^16^.

*Inclusion/exclusion criteria/ screening*

All children, who volunteered for the HUBU project, were included, except for those with any known history of neurological or psychiatric disorders or significant brain injury, according to parent reports. Nine HUBU participants were excluded from the present study because of incidental clinical findings on the MRI scans (n=1), receiving a clinical diagnosis later after inclusion (n=2), or because being a sibling of one of the other participants (n=6), reducing the sample from 95 to the above mentioned 86.

*Variables used for education, income and general cognitive ability (GCA)*

Highest level of maternal and paternal education was acquired for all participants in the 2^nd^ and 6^th^ assessments and translated into years of education using national norms. Average years of parental education, or years of education from one parent if unavailable for both parents, were used in the analysis. Furthermore, parental income (eight intervals: <13,400; 13,400-26,800; 26,800-40,200; 40,200-53,600; 53,600-67,000; 67,000-80,300; 80,300-93,700; 93,700< EUR) was assessed in the 6^th^ assessment from 65 of the included participants. The HUBU study did not include measures of general cognitive ability.

**LCBC**

*Population, recruitment and general description of study/ procedures*

Cognitively healthy, community dwelling participants across the lifespan were drawn from studies coordinated by the Research Group for Lifespan Changes in Brain and Cognition (LCBC [www.oslobrains.no](http://www.oslobrains.no)), approved by a Norwegian Regional Committee for Medical and Health Research Ethics. Written informed consent was obtained from all adult participants and from parents or other legal guardians for participants below age of majority. The samples were recruited by newspaper and web page adds, and part of the developmental sample was recruited through the population registry study MoBa <https://www.fhi.no/en/studies/moba/> Most participants, including all children, were recruited for observational studies, while some adults were recruited to enter into cognitive training studies after baseline assessment. For descriptive reasons, and to enable comparison of effects across samples of development and adulthood, the LCBC samples were divided by age at baseline assessments (included in the present study), so that *LCBC dev* comprise participants below 20 years of age, and *LCBC adult* comprise participants 20 years of age and above.

*Inclusion/exclusion criteria/ screening*

Adult participants were screened using a standardized health interview prior to inclusion in the study. Participants with a history of self- or parent-reported neurological or psychiatric conditions, including clinically significant stroke, serious head injury, untreated hypertension, diabetes, and use of psychoactive drugs within the last two years, were excluded. Further, participants reporting worries concerning their cognitive status, including memory function, were excluded. All participants above 40 yearsscored ≥24 on the Mini Mental State Examination ^17^.

*Variables used for education, income and general cognitive ability (GCA)*

Education was recorded as total years of education to the highest obtained degree, for LCBC-adult, and for LCBC-dev, average of parental education was used. Age was recorded in years and months as the mean of age at the time of MRI scan and the age at the time of the cognitive testing. Income was recoded as yearly nett income, NOK converted to € with the rate of 1NOK to 0.10€. For data prior to 2016, income was recoded in bins of 100k NOK starting from below 199k to 700 and above and means for each bin was used (100k, 350k, 450k, 550, 650k, 800k). Crystallized and Fluid GCA was recorded based on either the two-subtest (Vocabulary, Matrix reasoning) or 4 subtest (Vocabulary, Similarities, Matrix reasoning and Block design) version of the WASI ^13^ for participants above 6.5 years, and for children below 6.5, years, based on corresponding subtests from WPPSI-III ^18^.

**NESDA**

*Population, recruitment and general description of study/ procedures*

Participants were recruited from the Netherlands Study of Depression and Anxiety (NESDA), a large-scale, multisite, longitudinal, observational cohort study recruited from the general community, general practitioners and specialized mental health care centers (for details see ^19^). Of the 2981 NESDA respondents, a subgroup of participants was asked to participate in the NESDA neuroimaging study (n=301). Participants were eligible to participate in the neuroimaging study if they were aged between 18 and 57 years, met the DSM-IV criteria for diagnosis of major depressive disorder (MDD) and/or an anxiety disorder (panic disorder, social anxiety disorder, and/or generalized anxiety disorder) in the 6 months preceding the baseline interview, or had no lifetime DSM-IV diagnosis (i.e. healthy controls). NESDA neuroimaging exclusion criteria for patients were the presence of axis-I disorders other than MDD, panic disorder, social anxiety disorder, and/or generalized anxiety disorder and any use of psychotropic medication other than stable use of SSRIs or infrequent benzodiazepine use (i.e. equivalent to 2 doses of 10 mg of oxazepam 3 times per week or use within 48 hours prior to scanning). NESDA neuroimaging exclusion criteria for all participants were the presence or history of major internal or neurological disorders, dependence on or recent abuse (past year) of alcohol and/or drugs, hypertension, and general magnetic resonance imaging contraindications. Healthy controls were currently free of, and had never met criteria for, depressive or anxiety disorders or any other axis-I disorder and were not taking any psychotropic medication. Diagnoses according to DSM-IV algorithms were established using the structured Composite International Diagnostic Interview, lifetime version 2.1^20^, administered by a trained interviewer. Overall, 301 native Dutch-speaking participants (233 patients and 68 controls) were included and underwent magnetic resonance imaging in 1 of 3 participating centers: Leiden University Medical Center, Amsterdam Medical Center, and University Medical Center Groningen. Scanning took place between 2005 and 2007**.** Of these, 11 were excluded due to bad image quality and two did not complete the scan due to claustrophobia, reducing the final sample to 288. Age was entered as the age in years at time of baseline interview. Education was based on the self-reported highest attained degree, recoded to number of years of education according to the Dutch Central Bureau of Statistics. Income was based on self-reported net household income per month in euros as a 24-category variable, recoded to middle income per bin. The ethical review boards of each center approved this study. All participants provided written informed consent after receiving written information. NESDA did not include measures of general cognitive ability.

**UB**

*Population, recruitment and general description of study/ procedures, inclusion/ exclusion criteria/ screening, variables used for education, income and general cognitive ability (GCA), MRI scanning and processing*

Cohorts from the University of Barcelona (UB) site were collapsed across a number of substudies described below.

*WAHA cohort ^21^:* Recruitment and selection of participants took place between May 2012 and May 2014; the trial ended May 31, 2016. Participants were healthy elderly men and women with normal cognitive and visual function. Inclusion criteria were age between 63 and 79 years, apparently healthy, and equally willing to be in either of the two groups. Exclusion criteria included inability to undergo neuropsychological testing; morbid obesity (BMI ≥ 40 kg/m^2^); uncontrolled diabetes (HbA1c > 8%); uncontrolled hypertension (on-treatment blood pressure ≥ 150/100 mmHg); prior stroke, significant head trauma or brain surgery; relevant psychiatric illness; major depression; cognitive deterioration or dementia with a score < 24 on the Mini-Mental State Examination; other neurodegenerative disorders like Parkinson’s disease; advanced AMD or eye-related conditions precluding ophthalmological evaluation; prior chemotherapy; chronic illness with projected shortened lifespan; allergy to walnuts; customary use of fish oil and/or tree nuts (> 2 servings/week) and/or other relevant sources of ALA, such as flaxseed oil or soy lecithin. Eligible participants were recruited via mailing study brochures (LLU) or through the non-profit organization Institute of Aging (BCN), advertisements in the study centers, and word of mouth. Interested individuals attended an informational group meeting, completed a short medical questionnaire and signed the informed consent. Next candidates had a face-to-face interview with the study clinician, who assessed potential compliance, reviewed the medical history, inclusion and exclusion criteria, and recent blood work and use of medications or supplements, and administered the MMSE. Eligible participants were scheduled to have baseline tests (neuropsychological and ophthalmologic evaluations and collection of fasting blood and urine) and were then randomized to either the control or walnut group using a computerized random number table with stratification by center, sex, and age range. Couples entering the study were treated as one number and were randomized into the same group. Block design from WAIS-III ^22^ and National Adult Reading Test (NART) ^23^ were used for testing GCA.

*CR and iTBS cohorts ^24^:* Healthy volunteers older than 60 were recruited via the Institute of Aging, Barcelona. Individuals willing to participate were gathered in an informal meeting to tell them about the investigation, which included repetitive transcranial magnetic stimulation (TMS). Eligible participants had a normal cognitive profile with MMSE scores ≥ 24 and performances not below 1.5 SD according to normative scores (adjusted for age and education (Peña-Casanova et al., 2009)) on a neuropsychological evaluation that covered the major cognitive domains (including: Verbal memory: Rey auditory verbal learning test; visual memory: Rey-Osterrieth complex figure; Language: Benton naming test; semantic and phonetic fluencies; Frontal/Executive functions: direct and inverse digits, symbol digits modalities test, trail making test, Stroop test, London tower test; Visuospatial: line orientation, and visuoperceptive: Popplereuter’s embedded figures test). Vocabulary from WAIS-III ^22^ and NART ^23^ were used for testing GCA.

*GABA cohort ^25^:* Participants were recruited from the Institute of Aging (Barcelona) and the University of Experience, an initiative by the University of Barcelona for students aged 55 and older, offering special one and two-year degrees. Older adults, aged ≥ 60 years (age (mean ± *SD*), 68.15 ± 4.6 years; age range: 60–79 years; 22 females), naive to stimulation, participated in this study after giving informed consent, in accordance with the Declaration of Helsinki (1964, last revision 2013). None of the participants reported a diagnosis of a neurological or psychiatric disorder or any TMS contraindication. Inclusion criteria for the older subjects included a normal cognitive profile with mini-mental state examination (MMSE) scores of ≥ 24 and performance scores not more than 1.5 standard deviation (*SD*) below normative data (adjusted for age and years of education) on any of the administered neuropsychological tests (i.e., they did not fulfill the criteria for mild cognitive impairment (MCI). The neuropsychological battery included (1) a screening test for dementia, using the MMSE, and an evaluation of: (2) premorbid cognition and intelligence quotient (IQ), using the vocabulary subtest of the Wechsler Adult Intelligence Scale-III (WAIS-III) and National Adult Reading Test (NART); (3) verbal memory, using the Free and Cued Selective Reminding Test (SRT); (4) executive functions, using the phonemic fluency task and Trail Making Test B (TMTB); (5) language, using the semantic fluency task and Boston Naming Test (BNT); and (6) speed of processing, using the Symbol Digit Modalities Test (SDMT). Vocabulary from WAIS-III ^23^ and NART ^23^ were used for testing GCA.

*PD and MSA cohorts ^26^:* Healthy volunteers were recruited from the Aging Institute in Barcelona. Inclusion criteria were: scores within normality in a battery of neuropsychological tests described below (adjusted for age and education), IQ within normality as measured by the Vocabulary subtest (WAIS, scalar score >7). Exclusion criteria included: diagnosis of any neurological or psychiatric disease, mild cognitive impairment (MCI), any other condition that might affect cognition (chemotherapy or radiotherapy), diagnosis of fibromyalgia and any incompatibility to undergo MRI. Visuospatial and visuoperceptual functions were assessed with Benton Visual Form Discrimination (VFD) and Judgment of Line Orientation (JLO) tests; executive functions were evaluated with phonemic (words beginning with the letter “p” in 1 minute) and semantic (animals in 1 minute) fluencies; memory through total learning recall (sum of correct responses from trial I to trial V) and delayed recall (total recall after 20 min) through scores on Rey’s Auditory Verbal Learning Test (RAVLT). Attention and WM were assessed with Digit Span Forward and Backward, the Stroop Color-Word Test, Symbol Digits Modalities Tests (SDMT) and the Trail Making Test (in seconds), part A (TMTA) and part B (TMTB); and language was assessed by the total number of correct responses in the short version of the Boston Naming Test (BNT). For the PD cohort, vocabulary from WAIS-III ^22^ was used to test GCA, while for MSA similarities from WAIS-IV ^27^ and vocabulary from WAIS-III were used.

**Whitehall II Imaging sub-study**

*Population, recruitment and general description of study/ procedures*

The Whitehall II study, starting in 1985, includes 10.308 British civil servants followed over time. At study start, phase 1 (1985-1988), the population representative cohort was in the age range 35-55 years. A random sample of 800 participants were recruited at Phase 11 (2012-2013) to undergo MRI scanning and cognitive testing at the University of Oxford between 2012-2016, at which time the total number participants was 6035, and the age range was 60-85 years ^28^. This sub-cohort of the Whitehall study is included in the present analyses. Ethical approval was granted generically for the “Protocol for non-invasive magnetic resonance investigations in healthy volunteers” (MSD/IDREC/2010/P17.2) by the University of Oxford Central University/ Medical Science Division Interdisciplinary Research Ethics Committee (CUREC/MSD-IDREC), who also approved the specific protocol: “Predicting MRI abnormalities with longitudinal data of the Whitehall II sub-study” (MSD-IDREC-C1-2011-71).

*Inclusion/exclusion criteria/ screening*

A random selection of 800 participants from those giving consent to be contacted at Phase 11 of the Whitehall II study was made. Exclusion criteria were MRI contraindications, or being unable to travel to Oxford without assistance. After the exclusion of those with excess motion (N=4) and self-reported history of stroke (N=16), the final sample consisted of N = 780 with MRI, cognitive test data and information about education. No participant was diagnosed with dementia.

*Variables used for education, income and general cognitive ability (GCA)*

Age and years of education were defined at time of scan. Education years were calculated as the difference between the age at which the participant commenced primary school and the age at which they first left full-time education. Total household income was defined at Phase 5 (1997-1999) on the following scale: 1 = <£999, 2 = £1K - <£3K, 3 = £3K - <£5K, 4 = £5K - <£8K, 5 = £8K - <£10K, 6 = £10K - <£20K, 7 = £20K - <£40K, 8 = £40K - <£60K, 9 = £60K - <£100K, 10 = £100K - <£200K, 11 = >£200K. For computing the fluid GCA component for this sample, the Digit Coding (DC) test scores from the Wechsler Adult Intelligence Scale - Fourth Edition (WAIS-IV; ^27^) were used. In the DC test participants have to write the appropriate novel symbol for each number within a given time. For computing the crystallized GCA component, the Test of Premorbid Functioning (TOPF; ^29^) scores were used. The TOPF consists of a list of seventy written words, which must be read aloud and is marked according to pronunciation. The TOPF is used to estimate an individual's level of intellectual functioning before the onset of injury or illness. We calculated premorbid IQ from the raw score, adjusted for sex and years of education.

**ABCD**

*Population, recruitment and general description of study/ procedures*

The primary aim of the Adolescent Brain Cognitive Development (ABCD) study is to track human brain development from childhood through adolescence to determine biological and environmental factors that impact or alter developmental trajectories ^30^. ABCD has recruited > 10 000 9-10 years olds across 21 US sites with harmonized measures and procedures, including imaging acquisition <https://abcdstudy.org/scientists-workgroups.html>. A goal of the ABCD study is that its sample should reflect, as best as possible, the sociodemographic variation of the US population ^31^. For ABCD, the dataset release 2.0.1 was used, consisting of 11875 participants at baseline.

*Inclusion/exclusion criteria/ screening*

2105 twins and 30 triplets were excluded, leaving 9740 participants, of whom 9049 had underwent MRI scanning with FreeSurfer Quality Control OK.

*Variables used for education, income and general cognitive ability (GCA)*

Education (ABCD data-field high.educ) was entered as the highest education of parents (1: < HS Diploma, 2:HS Diploma/GED, 3: some college, 4: Bachelor, 5: Post Graduate Degree). Income (ABCD data-field demo_comb_income_p) was entered as the parents combined income, recoded to the middle of each income category and converted to Euros using the average USDxEuro exchange rate for 2019 according to US Internal revenue services (0.893). Behavior tests were taken from the NIH Toolbox (<http://www.healthmeasures.net/explore-measurement-systems/nih-toolbox>). Fluid intelligence score (ABCD data-field nihtbx_fluidcomp_uncorrected) was derived by averaging the normalized scores of each of the Toolbox tests that are fluid ability measures (Flanker, Dimensional Change Card Sort, Picture Sequence Memory, List Sorting and Pattern Comparison), then deriving scale scores based on this new distribution. The Crystallized intelligence score (ABCD data-field nihtbx_cryst_uncorrected) was derived by averaging the normalized scores of each of the Toolbox tests that are crystallized measures (Picture Vocabulary and Reading Tests), then deriving scale scores based on this new distribution.

*MRI scanning and processing*

Brain imaging data were collected across 21 sites on 3T scanners (Siemens Prisma (Siemens Corp., Erlanger, Germany), GE Discovery MR750 (GE Healthcare, Chicago, IL), and Philips Achieva (Philips, Amsterdam, the Netherlands). Aquisition parameters are listed in ^30^ and at <https://abcdstudy.org/images/Protocol_Imaging_Sequences.pdf>. Imaging preprocessing steps included gradient nonlinearity distortion correction, intensity inhomogeneity correction, and registration and resampling to a custom atlas brain with 1.0 mm isotropic voxels. Brain volumes were computed from the preprocessed T1 images using FreeSurfer version 5.3.0.

**HCP**

*Population, recruitment and general description of study/ procedures*

The US National Institute of Health (NIH) announced a Request for Applications for the Human Connectome Project (HCP) in 2009, with an overarching objective of studying human brain connectivity and its variability in healthy adults. In 2010, grants were awarded to two consortia (<http://www.neuroscienceblueprint.nih.gov/connectome/>). One is a 5-year grant to a consortium of ten institutions in the United States and Europe, led by Washington University and the University of Minnesota (the ‘WU-Minn HCP Consortium’). This consortium aims to study brain connectivity and function with a genetically informative design in 1,200 individuals using four MR-based modalities plus MEG and EEG. Behavioral and genetic data are also acquired from these participants. The HCP aimed to study and freely share data from 1200 young adults (ages 22-35) from families with twins and non-twin siblings, using a protocol that includes structural and functional magnetic resonance imaging (MRI, fMRI), diffusion tensor imaging (dMRI) at 3 Tesla (3T) and behavioral and genetic testing. The dataset used was the 1200 Subjects Release (<https://www.humanconnectome.org/storage/app/media/documentation/s1200/HCP_S1200_Release_Reference_Manual.pdf>). Age was entered as years without decimals.

*Inclusion/exclusion criteria/ screening*

The participant pool comes from healthy individuals born in Missouri to families that include twins, based on data from the Missouri Department of Health and Senior Services Bureau of Vital Records. 243 MZ or DZ twin pairs identified by genotyping were excluded from the present analyses, reducing the total number of participants from 1206 to 720. Additional recruiting efforts in HCP were used to ensure that participants broadly reflect the ethnic and racial composition of the U.S. population as represented in the 2000 decennial census. ‘Healthy’ was broadly defined, aiming for a pool that is generally representative of the population at large, to capture a wide range of variability in healthy individuals with respect to behavioral, ethnic, and socioeconomic diversity. Sibships with individuals having severe neurodevelopmental disorders (e.g., autism), documented neuropsychiatric disorders (e.g., schizophrenia or depression) or neurologic disorders (e.g., Parkinson's disease) were excluded. Individuals with illnesses such as diabetes or high blood pressure were additionally excluded, as were non-twins (current sample) born prior to 37 weeks gestation. Individuals who were smokers, overweight, or had a history of heavy drinking or recreational drug use without having experienced severe symptoms were included. For a full list of inclusion and exclusion criteria, see ^32^. This reduced the HCP sample for the current analyses to 589, of whom 538 had MRI.

*Variables used for education, income and general cognitive ability (GCA)*

Education (HCP data-field SSAGA_Educ) was coded as years of education completed: <11 = 11; 12; 13; 14; 15; 16; 17+ = 17. Income (HCP data-field SSAGA_Income) was recoded to the middle of each income category and converted to euros using the average USDxEuro exchange rate for 2019 according to US Internal revenue services (0.893). Behavior tests were taken from the NIH Toolbox (<http://www.healthmeasures.net/explore-measurement-systems/nih-toolbox>). Fluid intelligence score (HCP data-field CogFluidComp_Unadj) was derived by averaging the normalized scores of each of the Toolbox tests that are fluid ability measures (Flanker, Dimensional Change Card Sort, Picture Sequence Memory, List Sorting and Pattern Comparison), then deriving scale scores based on this new distribution. The Crystallized intelligence score (HCP data-field CogCrystalComp_Unadj) was derived by averaging the normalized scores of each of the Toolbox tests that are crystallized measures (Picture Vocabulary and Reading Tests), then deriving scale scores based on this new distribution. The participant scores for the cognitive categories above were normed to those in the entire NIH Toolbox Normative Sample (18 and older), regardless of age or any other variable, where a score of 100 indicates performance that was at the national average and a score of 115 or 85, indicates performance 1 SD above or below the national average.

*MRI scanning and processing*

Imaging data were collected and processed by the human connectome project (https://www.humanconnectome.org/study/hcp-young-adult) as described in ^33^. Imaging data were collected a customized Siemens 3T “Connectome Skyra” housed at Washington University in St. Louis, using a standard 32-channel Siemens receive head coil and a “body” transmission coil designed by Siemens specifically for the smaller space available using the special gradients of the WU-Minn and MGH-UCLA Connectome scanners. Anatomical T1-weighted magnetization-prepared rapid gradient echo (MPRAGE) images were obtained in the sagittal plane at 0.7mm isotropic resolution, and T2 weighted FLAIR images were acquired at identical resolution in the sagittal plane. Images were processed using a custom combination of tools from FSL and FreeSurfer ^34-36^. FreeSurfer estimates of intracranial and gray and matter volume (sum of bilateral segmentations) were entered for analysis (HCP data-fields FS_IntraCranial_Vol, FS_Total_GM_Vol).

**UKB**

*Population, recruitment and general description of study/ procedures*

UK Biobank (UKB) (<https://www.ukbiobank.ac.uk/about-biobank-uk/>) is a major national and international health resource with the aim of improving the prevention, diagnosis and treatment of a wide range of illnesses. UK Biobank recruited ≈500,000 people aged between 40-69 years in 2006-2010 from across the country to take part in this project ^37^. Potential participants were identified through National Health Service (NHS) registers according to being aged 40-69 and living within a reasonable travelling distance of an assessment centre. Assessment centres (22 in total) are located in accessible and convenient locations with a large surrounding population. Participants have undergone measures and provided samples and detailed information about themselves and agreed to have their health followed. The dataset released February 2020 was used. Age was calculated from year and month of birth (day of month is missing, and was set to 1 for all subjects) to date of assessment. Age was calculated at the 0_0 timepoint for participants without MRI, and 2_0 timepoint for participants with MRI. Behavior data were collected at the 0_0 timepoint for participants without MRI, and at the 2_0 timepoint for participants with MRI (i.e. at the same timepoint as the MRI data collection).

*Inclusion/exclusion criteria/ screening*

11244 participants (6215 female) reported being part of a multiple birth and were excluded from the analyses.

*Variables used for education, income and general cognitive ability (GCA)*

Education: For the Biobank participants` generation, the UK school system provided free universal compulsory education between the ages of 5 and 15 to 16 years. Highest educational qualification was categorized into 4 levels, as in ^37^, based on the UK Biobank categories coded in field 6138 (1: None, 2: O-levels or Certificate of Secondary Education or equivalent, 3: A-levels/AS-levels, National Vocational Qualification, Higher National Diploma, or Higher National Certificate or equivalent, or other professional qualifications eg: nursing, teaching, 4: College or University degree, See <http://www.ukbiobankeyeconsortium.org.uk/sites/www.ukbiobankeyeconsortium.org.uk/files/Role%20of%20Educational%20Exposure%20in%20the%20Association%20Between%20Myopia%20and%20Birth%20Order.pdf>). Income was entered as annual household income (UKB Data-field 738) at time of examination. Income was entered as the middle of each income category and converted to euros using the middle UK pounds to Euro exchange rate for 2019 according to Bank of England (1.1405). General cognitive ability was entered as the raw fluid intelligence score (UKB Data-field 20016). This is a simple unweighted sum of the number of correct answers given to the 13 fluid intelligence questions.

*Genetic Ancestry Factor (GAFs)*

We estimated the Genetic Ancestry Factor for the UKB sample and the LCBC developmental and adult sample. Since these samples have been genotyped by different arrays we estimated these factors separately using the program PLIKN (v1.9)^38^. The estimated factors have been used as covariates in our models to capture unmodeled ancestral difference among participants.

LCBC sample: The sample included in the present study largely overlaps with the one reported by Walhovd et al. ^39^. Briefly, DNA was extracted from either saliva or buccal swabs in Oslo or from the LIGA laboratory at University of Lübeck, Germany. Extracted DNA samples were quantified, quality controlled, normalized and aliquoted at LIGA. The Global Screening Array (GSA; Illumina, Inc.) was used to genotype the DNA sample. Genotype calling was performed by GenomeStudio v2.0.4. In total 696,375 variants were obtained. These called variants and individuals were further quality controlled by the PLINK program^38^ to remove variants that showed large missingness, or low frequencies, or violated the Hardy Weinberg Equilibrium, and to remove subjects with ambiguous predicted sex, or abnormal heterozygosity, or low call-rate, and to remove variants that showed large missingness, or low frequencies, or violated the Hardy Weinberg Equilibrium, from further analysis. The genetic ethnicity for remaining participants was estimated using the quality controlled genotypes (further details see Walhovd et al.^39^). First, these genotypes were merged with the genotypes from 1000 Genomes Project phase 3 superpopulations; second, the –pca command from the PLINK program was used to estimates the principal components for the merge dataset. The package class (v2.3.2) from the software R was used to identify non-European participants, which have been excluded in the present study. The remaining genetically determined European samples were used to compute genetic ancestry factors used in this study. Only SNPs that have a minor allele frequency (MAF) >= 0.1 were used. The program PLINK was first used to obtain nearly independent SNP set using parameters, --indep-pairwise 100 50 0.1. Then, the remaining SNPs were included in the GAF estimation with the PLINK command, --pca. The top 10 principal components were taken as GAFs used as covariates in the statistical analyses.

UK biobank sample: The UK biobank sample was genotyped by the UK Biobank Axiom array from Affymetrix or the UK BilEVE array. We only used participants that showed a kinship coefficient 0 (provided by the UK biobank). In addition, participants that failed the UKBiLEVE genotyping, or are outliers of heterozygosity estimates, or having high missingness were excluded from this analysis. The quality controls on variants has been performed by UK biobank and we did not perform further quality controls(for further details please see <https://www.ukbiobank.ac.uk/scientists-3/genetic-data/>). We computed the GAF for the remaining data based on SNPs that have MAF > 0.1 and are nearly independent. The same command used for LCBC sample was used for UKB sample. The top 10 principal components were taken as GAFs in our statistical analysis.

*Ethnicity*

The codes used for ethnicity and the distributions are shown below:

Ethnicity in UKB (field 1001)

White -> white

Mixed -> other_mixed

Asian -> asian

Black -> black

Chinese -> asian

Other ethnic group -> other_mixed

Distribution:

white	461992

asian	11320

black	7849

other_mixed	7364

Ethnicity in ABCD

White -> white

Black -> black

Hispanic -> hispanic

Asian -> asian

Other -> other_mixed

Distribution:

white	4801

hispanic	2163

black	1486

other_mixed	1033

asian	243

Missing	14

Ethnicity in HCP:

White -> white:

Black or African Am. -> black

Asian/Nat. Hawaiian/Othr Pacific Is. -> Asian

More than one -> other_mixed

Am. Indian/Alaskan Nat. -> other_mixed

Distribution:

white	388

black	111

asian	47

other_mixed	24

Missing	19

*MRI scanning and processing*

Imaging data were collected and processed by the UK Biobank (https://www.ukbiobank.ac.uk) as described in ^40^. Imaging data were collected using 3.0 T Siemens Skyra (32-channel head coil). Anatomical T1-weighted magnetization-prepared rapid gradient echo (MPRAGE) images were obtained in the sagittal plane at 1mm isotropic resolution, and T2 weighted FLAIR images were acquired at 1.05x1x1mm resolution in the sagittal plane. Images were processed by the UK Biobank using the FreeSurfer 6.0 software package.

**Specific information on image acquisition in the different samples**

T1 weighted structural scans were acquired at Siemens, Philips and GE scanners at the various sites. The sequence parameters for each cohort are given in the Supplementary Table 1.

| **Sample** | **Scanner** | **Tesla** | **Sequence parameters** |
| --- | --- | --- | --- |
| BASE-II | Tim Trio Siemens | 3.0 | TR: 2500 ms, TE: 4.77 ms, TI: 1100 ms, flip angle: 7°, slice thickness: 1.0 mm, FoV 256×256 mm, 176 slices |
| Betula | Discovery GE | 3.0 | TR: 8.2 ms, TE: 3.2 ms, TI: 450 ms, flip angle: 12°, slice thickness: 1 mm, FOV 250×250 mm, 176 slices |
| CALM | Tim Trio  Siemens | 3.0 | TR: 2250 ms, TE: 3.02 ms, TI: 900 ms, flip angle: 9°, slice thickness 1 mm, 192 slices, GRAPPA = 2 |
| Cam-CAN | Tim Trio  Siemens | 3.0 | TR: 2250 ms, TE: 2.98 ms, TI: 900 ms, flip angle: 9°, slice thickness 1 mm, FOV 256×240 mm, 192 slices |
| HUBU | Tim Trio Siemens | 3.0 | TR: 1550 ms, TE: 3.04 ms, TI: 800 ms, flip angle: 9°, slice thickness 1 mm, FOV 256×256 mm, 192 slices |
| LCBC | Avanto Siemens | 1.5 | TR: 2400 ms, TE: 3.61 ms, TI: 1000 ms, flip angle: 8°, slice thickness: 1.2 mm, FoV: 240×240 m, 160 slices, iPat = 2 |
|  | Avanto Siemens | 1.5 | TR: 2400 ms, TE = 3.79 ms, TI = 1000 ms, flip angle = 8, slice thickness: 1.2 mm, FoV: 240 x 240 mm, 160 slices |
|  | Skyra Siemens | 3.0 | TR: 2300 ms, TE: 2.98 ms, TI: 850 ms, flip angle: 8°, slice thickness: 1 mm, FoV: 256×256 mm, 176 slices |
|  | Prisma Siemens | 3.0 | TR: 2400 ms, TE: 2.22 ms, TI: 1000 ms, flip angle: 8°, slice thickness: 0.8 mm, FoV: 240×256 mm, 208 slices, iPat = 2 |
| NESDA | Intera  Phillips | 3.0 | TsssR: 9ms, TE 3.5ms, prepulse delay: 1000 ms, flip angle: 8°, slice thickness: 1mm, FoV: 256x 256, 170 slices |
| UB | Tim Trio Siemens | 3.0 | TR: 2300 ms, TE: 2.98, TI: 900 ms, slice thickness 1 mm, flip angle: 9°, FoV 256×256 mm, 240 slices |
| WH-II | Verio Siemens | 3.0 | TR: 2530 ms, TE: 1.79/3.65/5.51/7.37 ms, TI: 1380 ms, flip angle: 7°, slice thickness: 1.0 mm, FOV: 256×256 mm |
| ABCD | Prisma  Siemens | 3.0 | TR: 2500 ms, TE: 2.88 ms, TI: 1060 ms, flip angle: 8°, slice thickness: 1 mm, FoV: 256×256 mm, 176 slices, parallel imaging = 2 |
|  | Achieva/  dStream/  Ingenia  Phillips | 3.0 | TR: 6.31 ms, TE: 2.9 ms, TI: 1060 ms, flip angle: 8°, slice thickness: 1 mm, FoV: 256×240 mm, 225 slices, parallel imaging = 1.5 x 2.2 |
|  | MR750/  DV25-26  GE | 3.0 | TR: 2500 ms, TE: 2 ms, TI: 1060 ms, flip angle: 8°, slice thickness: 1 mm, FoV: 256×256 mm, 208 slices, parallel imaging = 2x |
| HCP | Connectome  Skyra  Siemens* | 3.0 | TR: 2400 ms, TE: 2.14 ms, TI: 1000 ms, flip angle: 8°, slice thickness: 0.7 mm, FOV: 224 mm, 256 slices, GRAPPA = 2 |
| UKB | Skyra  Siemens | 3.0 | TR: 2000 ms, TI: 880 ms, slice thickness: 1 mm, FoV: 208×256 mm, 256 slices, iPAT=2 |

**Supplementary Table 2** MR acquisition parameters

TR: Repetition time, TE: Echo time, TI: Inversion time, FoV: Field of View, iPat: in-plane acceleration, GRAPPA: GRAPPA acceleration factor. *Customized

**Statistical analyses**

For all measures of neuroanatomical volume (GM and ICV), we ran separate regressions for each cohort and each neuroanatomical measure predicting volume by age and checking whether absolute residuals exceeded a four standard-deviations criterion. If so, the respective participants were entirely deleted from the analysis. In total, we deleted 26 participants from the analysis. We excluded 1 participant from BASE-II, 1 participant from LCBC-adult, 3 participants fromLCBC-Dev, 18 participants from UKB, and 3 participants from ABCD.
